# Kv1.3 inhibition alleviates neuropathology via neuroinflammatory and resilience pathways in a mouse model of Aβ pathology

**DOI:** 10.64898/2025.12.25.696456

**Authors:** Rashmi Kumari, Christine Bowen, Upasna Srivastava, Amanda Dabdab Brandelli, Prateek Kumar, Dilpreet Kour, Sneha Malepati, Wooyoung Eric Jang, Mark Bromwich, Hollis Zeng, Anson Sing, Steven A. Sloan, Heike Wulff, Srikant Rangaraju

**Author notes:** Corresponding author Rashmi Kumari, Srikant Rangaraju. Equally contributed as first author (Rashmi Kumari, Christine Bowen, Upasna Srivastava).

## Abstract

Inhibition of voltage-gated potassium channel Kv1.3 is a therapeutic strategy to curb microglia-mediated neuroinflammation in neurodegeneration, although the cellular and signaling mechanisms of disease-modification by Kv1.3 blockers are unclear. In this study, we delineate protective mechanisms of Kv1.3 blockade in a mouse model of Alzheimer’s disease (AD) pathology using comprehensive transcriptomics and proteomics profiling of brain, corresponding with neuropathological effects of two translationally relevant Kv1.3 blockers, namely small molecule PAP-1 and peptide ShK-223. Following 3 months of treatment, both molecules reduced Ab plaque burden. Single nuclear RNA seq (snRNA seq) of brain nuclei showed that PAP-1 disproportionately impacted oligodendrocytes and microglia and increased crosstalk between neurons and astrocytes with endothelial cells. In contrast, ShK-223 had pronounced effects on glutamatergic neurons and astrocytes. Both blockers increased expression of myelination genes in oligodendrocytes and synaptic genes in neurons. Neuroprotective effects of PAP-1 were further confirmed by bulk brain transcriptomics and proteomics whereby PAP-1 increased levels of synaptic, cognitive resilience and mitochondrial proteins, while decreasing glial and immune pathways including STAT1/3 phosphorylation. Using proximity labeling and co-immunoprecipitation, we found that Kv1.3 interacts with STAT1/3 in microglia. Using microglial cell lines and primary microglia, we discovered a preferential functional coupling between Kv1.3 and type 2 but not type 1 IFN signaling. Brain-level disease modification by Kv1.3 blockade was reflected in the cerebrospinal fluid (CSF) via reduced levels of neurofilament-light (NEFL) and resilience protein RPH3A, both of which are increased in human AD CSF. Together, this study demonstrates functional links between Kv1.3 channels and type 2 IFN signaling and reveals distinct cellular effects of Kv1.3 blockers in AD pathology that correspond with reduced neuropathology and neuroinflammation, augmentation of resilience and neuro-vascular pathways, along with biomarkers of therapeutic effect.

## INTRODUCTION

Alzheimer’s disease (AD), one of the most common neurodegenerative diseases, is a multi-cellular disorder in which the cumulative effects of protein aggregation, dysfunction in neurons, glial cells and cerebrovascular components result in several pathological alterations in the brain like synaptic loss, neuronal dystrophy, gliosis and associated vascular changes^1^. Neuropathologically, AD is characterized by aggregation of amyloid beta (Aβ) as well as neurofibrillary tangles which can activate pathogen recognition receptors (PRRs) triggering the release of several inflammatory mediators. Microglia and astrocytes are the major cell types involved in these neuroinflammatory cascades^2^. Astrocytes play a key role in metabolism, synapse formation, and regulation in the cellular phase of disease^3^. Reactive astrogliosis, a complex and dynamic response of astrocytes marked by proliferation, hypertrophy, and increased expression of several intermediate filaments, is an early event in both mouse models of AD and patients with prodromal AD^4, 5^.

Microglia, the major enactors of neuroinflammation in the brain, show intricate associations with senile plaques in AD and play a critical role in phagocytosis and clearance of cellular debris and protein aggregates along with synaptic pruning^6, 7, 8^. The technical advancement of transcriptional profiling through single cell RNA seq and spatial transcriptomics further reveals microglial heterogeneity and suggests existence of different microglial populations that exhibit distinct genetic and functional profiles, majority of which are conserved in health and disease conditions^9, 10, 11^. Microglial subtypes consisting of Disease Associated Microglia (DAMs) have been identified at different stages of disease in models of AD pathology as well as in human AD^12, 13, 14, 15^. Along with DAMs, a population of interferon responsive microglia (IRMs) characterized by increased expression of interferon stimulated genes has also been reported in these disease models^16^. Based on disease context and pathological stage, DAMs further exhibit functional heterogeneity and can either be neurotoxic or neuroprotective^14, 17^. The neurotoxic state is a chronic proinflammatory state of microglia responsible for increased production of Reactive Oxygen Species (ROS) and secretion of proinflammatory cytokines along with dysfunctional lysosomal deposits resulting in direct neuronal damage and impaired pathological protein clearance^18^. Regulators of these proinflammatory DAMs can serve as potential therapeutic targets in different neurological diseases, hence their identification and characterization are important. Recently, the potassium channel Kv1.3 has emerged as one of the regulators of proinflammatory DAMs with increased expression levels in both mouse models as well as acutely isolated microglia from human AD brain^19, 20, 21^.

Kv1.3 is a member of the *Shaker* family of voltage-gated ion channels responsible for potassium (K+) efflux upon membrane depolarization^22, 23, 24^. Several non-excitable cells like microglia, T-cell, and T effector memory cells (TEM) express this channel on the cell surface as well as mitochondria. K+ efflux via Kv1.3 is functionally coupled with membrane depolarization that can be induced by Ca^2+^ flux^25, 26^.

Recently, Kv1.3 has also been shown to be involved in different signaling pathways through protein-protein interactions, possibly by formation of macromolecular complexes^27, 28, 29^. While Kv1.3 has been primarily studied in microglia and T lymphocytes in neurological disorders, Kv1.3 mRNA and functional channels have been reported in other cell types including neurons, astrocytes, oligodendrocytes, and oligodendrocyte progenitor cells (OPCs)^30, 31, 32, 33^. Microglial Kv1.3 has been associated with neuroinflammation in different neuropathological conditions like Parkinson’s disease (PD), epilepsy, stroke, and AD^34^. Interestingly, upregulation of Kv1.3 transcript and protein has also been reported in other glial cells like astrocytes in a mouse model of Multiple Sclerosis^35^. The presence of Kv1.3 in cell types beyond microglia and T cells raises the possibility that translationally relevant Kv1.3 blockers may also impact the function of these cell types.

Both cell membrane permeant small molecules (eg. PAP-1) and membrane-impermeant peptide blockers of Kv1.3 (eg. ShK-related peptides) have been tested in preclinical models with their beneficial effects demonstrated in mouse models of AD, PD, epilepsy, and ischemic stroke^36, 37, 38, 39, 40^. However, the molecular mechanisms of neuroprotection and the effects of these pharmacological inhibitors on different brain cell types have not been explored so far. Considering the translational potential of Kv1.3 inhibitors, a comprehensive understanding of their cellular and molecular targets is crucial to defining mechanisms of neuroprotection to maximize their therapeutic efficacy. Given the relationship between pro-inflammatory immune pathways and Kv1.3 channel function in microglia, it is important to define immune signaling mechanisms that are regulated by Kv1.3 channels in immune cells. Lastly, to facilitate translation of pre-clinical findings to patients, it is also important to identify disease-relevant biofluid biomarkers that may reflect therapeutic efficacy of Kv1.3 blockers.

In this manuscript, we show that Kv1.3 blockade by two selective molecules (PAP-1 and ShK-223) alleviated disease pathology in an amyloidosis mouse model of AD (5xFAD), re-affirming prior findings ^21, 36^. Using single nuclear transcriptomics of brain, we observed differential cellular responses to both Kv1.3 blockers. PAP-1 preferentially impacted microglia and oligodendrocytes while ShK-223 preferentially impacted neurons and oligodendrocytes. We also observed shared effects of Kv1.3 blockers on ferroptosis and cell survival pathways in microglia. Kv1.3 blockade increased synaptic genes in both excitatory and inhibitory neurons consistent with a neuroprotective effect, along with augmentation of proteins that determine cognitive resilience (*e.g.,* NRN1). We further confirmed that Kv1.3 channels interact with STAT1/3 proteins in microglia and functionally regulate STAT signaling. Importantly, we observed that Kv1.3 channels preferentially regulate Type 2 IFN (gamma) signaling but not Type 1 IFN (alpha/beta) signaling. Consistent with these in vitro findings, Kv1.3 blockade reduced activation of STAT3 in 5xFAD brain, together nominating Kv1.3-mediated regulation of IFN-STAT signaling as a pathway for disease-modification by Kv1.3 blockers. Using cerebrospinal fluid from mice treated with Kv1.3 blockers, we also identified CSF proteomic biomarkers of therapeutic efficacy of Kv1.3 blockers, many of which are of relevance to human AD.

## RESULTS

### Kv1.3 inhibition by PAP-1 or ShK-223 reduces Aβ pathology in 5xFAD mice

We used 5xFAD mice, a widely used model of rapid and progressive Aβ pathology, to assess the pathological changes during disease progression. Previous studies report increased Aβ plaque pathology and neuroinflammation in both human AD patients and AD mice models with increased proinflammatory activation of microglia in female mice compared to males^21, 41, 42^. As expected, bulk proteomic MS data using an existing ageing cohort study with 10-month-old 5xFAD mice showed increased presence of Aβ in female mice compared to their age matched male controls **(Fig. 1A).** Next, we looked at the abundance of proteins of several markers of gliosis, synaptic integrity and neuroinflammation including GFAP, IBA, SV2A and STAT1/3 (**Supp. Fig 1A**) and further validated the presence of these proteins along with the phospho-forms of STAT1/3 in the brain lysates from the same cohort **(Fig. 1B & 1C Supp. Fig. 1D).** Both the protein abundance by MS and western blots showed increased markers of gliosis, neuroinflammation and a decrease in synaptic markers, specifically in the female mice. Given this sex-difference, we designed our Kv1.3 blockade study using the small molecule PAP-1 or the peptide ShK-223 in female 5xFAD mice.

**Figure 1.**
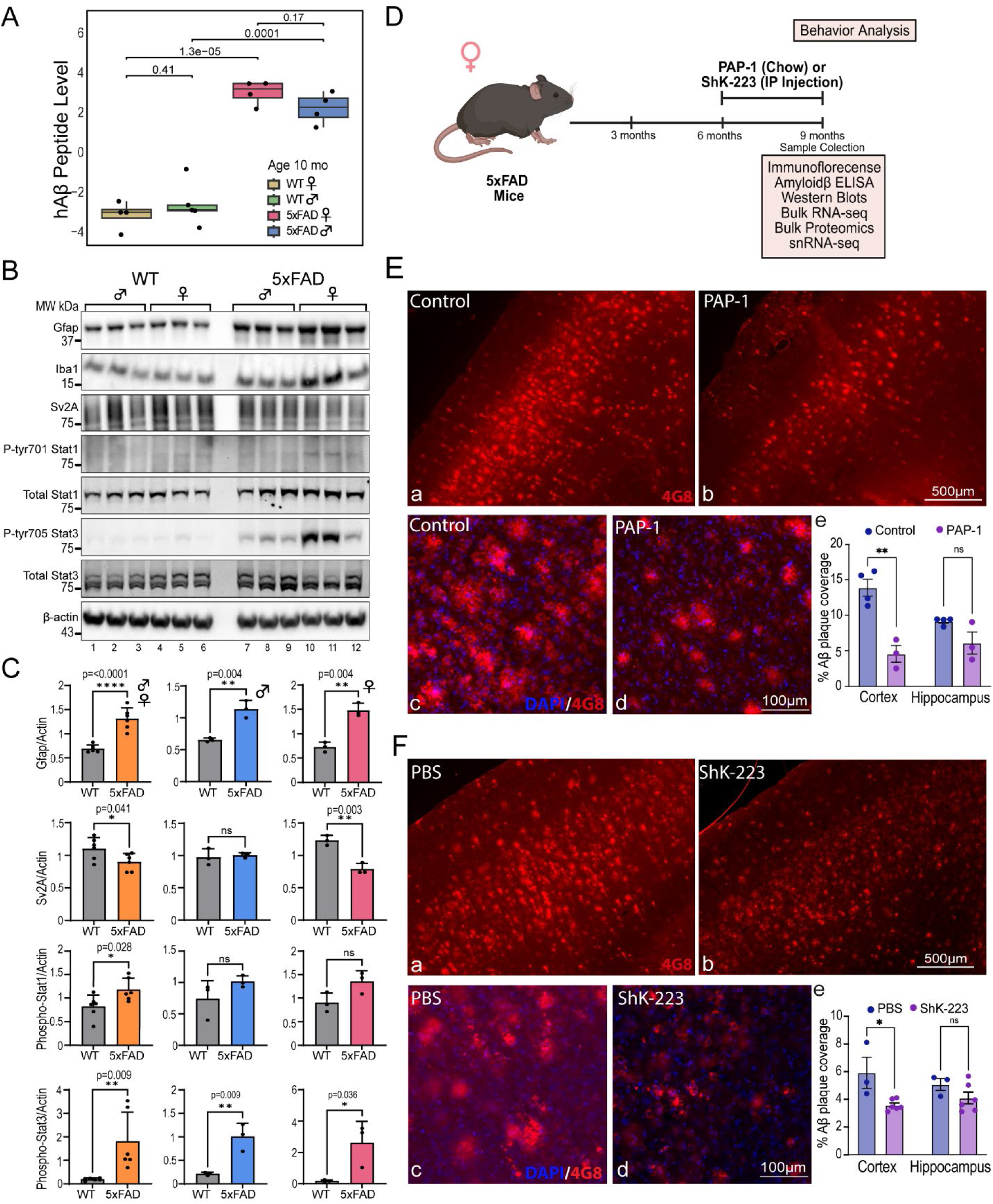
Blockade of Kv1.3 by PAP-1 or ShK-223 reduces Aβ pathology in 5xFAD mouse brain. **(A)** Protein abundance of Aβ from 10-month-old WT and 5xFAD mice (Male/Female) using tandem mass tag mass spectrometry. **(B & C)**Western blot and densitometric analysis of markers of synaptic integrity, gliosis and neuroinflammation. **(D)** Treatment paradigm for Kv1.3 blockade in 5xFAD mice, 6-month 5xFAD female mice were treated with either ShK-223 or PAP-1 for three months. Behavioral assessments (fear conditioning, Morris water maze) were performed to evaluate changes in learning and memory. Western blots, snRNA seq, proteomics, and IF were utilized to determine functional and biochemical changes. **(E & F)** IF images show a reduction in Aβ plaque load by PAP-1 and ShK-223 treatment. Mouse brain sections were stained for Aβ (n=3, 3 sections per mouse) and imaged at 4x for quantification and 20x for illustration using the Keyence. Aβ intensity is reduced in both PAP-1 and ShK-223 treated mice. Quantification of both PAP-1 and ShK-223 treated IF images show a statistical reduction of Aβ plaque counts in cortex and not hippocampus (unpaired T-test used). * p<0.05, ** p < 0.01, *** p< 0.001. See Supp Fig 1 for related supplemental analyses and figures.

Both these Kv1.3 blockers work differently and have different CNS bio-availabilities. The peptide ShK-223 binds to the external pore of the Kv1.3 channel, while PAP-1 binds to Kv1.3 on the intracellular side, leading to inhibition of K+ efflux^43, 44^ **(Supp. Fig. 1B**). Previous studies have shown that Kv1.3 blockade by PAP-1 or ShK-223 reduces Aβ plaque burden and neuroinflammation in Ab pathology models ^19, 21, 36^. We aged female 5xFAD mice to 6 months and treated them with PAP-1 (chow, 1100 ppm) or ShK-223 (2x a week i.p, 100 μg/kg/dose) for three more months (**Fig. 1D**). Following treatment, behavioral studies were performed on the 9 months old mice. Fear conditioning showed that treatment with PAP-1 and not ShK-223 resulted in a quicker training association of the context (shock) with the cue (light). This indicates that treatment with PAP-1 may increase the learning capacity of the 5xFAD mice (**Supp.Fig 1F**). Morris water maze showed no statistical differences between treatment and controls (**Supp.Fig. 1G**). Brain tissues were then harvested to evaluate biochemical, pathological, and cellular changes associated with Kv1.3 blockade. Immunofluorescence (IF) microscopy of sagittal cut tissues of 9 months old 5xFAD mice showed that Aβ plaque burden was reduced in the cortex by both ShK-223 and PAP-1 treatment. Both inhibitors induced a significant reduction of plaque counts only in the cortex but not the hippocampus which can be explained by the general sparsity of plaques in the hippocampus (**Fig. 1E & 1F)**(**Supp. Fig. 1E**). To further evaluate Aβ abundance, ELISA was preformed to determine total Aβ abundance. PAP-1 treatment reduced Aβ42 levels at 9 months, whereas ShK-223 treatment did not reduce total Aβ42 levels in brain homogenates (**Supp. Fig. 1C**). This suggests that PAP-1 reduces plaque and total levels of Aβ42 potentially by impacting both production and clearance mechanisms while ShK-223 only impacts plaque clearance mechanisms. These findings highlight that Kv1.3 inhibitors with different mechanisms of channel inhibition affect plaque clearance and Aβ42 levels differently.

### snRNA seq brain transcriptome profiling shows preferential cellular effects of PAP-1 and ShK-223

To examine brain cell type specific transcriptional responses to both PAP-1 and ShK-223 in these 5xFAD mice post Kv1.3 blockade, we performed snRNA seq on brain cortex-derived nuclei from 9 month old-5xFAD (n= 3 per group, PAP-1, control chow, i.p ShK-223, i.p saline) and WT mice (n=2, no treatment). We obtained single nuclear transcriptomes from 137,375 high quality nuclei that were classified into 24 clusters, that were further mapped to 7 parent cell types, based on published cellular markers **(Fig. 2A & Supp. Fig. 2A)**^45, 46^. The 24 clusters included 9 excitatory neurons (65%), 6 inhibitory neurons (14%), 1 oligodendrocyte (11%), 3 astrocyte (3.2%), 2 microglia (3%), 1 oligodendrocyte progenitor (2%), and 1 endothelial cluster (0.069%). We observed no differences in proportions of cell types across treatment groups in 5xFAD mice **(Supp. Fig. 2B).** Pseudobulk differential expression analyses were then performed to examine cell type-specific molecular changes. As compared to WT mice, microglia exhibited increased expression of DAM genes (*Hif1a, Tspo, Cd274, Csf1, Slc2a1*) and down-regulation of homeostatic genes (*Tmem119, Mfng, Stab1, Tspan18, Fgd2, Adrb2*) consistent with expected findings in the 5xFAD brain. We then assessed the effect of PAP-1 and ShK-223 treatment on distinct brain cell types, based on the proportion of DEGs within each population, because of Kv1.3 blockade, within each cell type. Interestingly, oligodendrocytes (28% DEGs) and microglia (26% DEGs) were the most responsive cell type upon PAP-1 treatment while ShK-223 treatment had a relatively stronger effect on glutamatergic neurons (21%) and oligodendrocytes (15%) as the major cell types with transcriptional alterations **(Fig. 2B & 2C).** These preferential effects on cell types cannot be explained by depth of transcriptomes within each cell type because there was no correlation between proportion of DEGs with transcriptome depth (**Supp. Fig. 2C**). Absolute number of DEGs, on the other hand, were highest in neuronal groups in both PAP-1 and ShK-223 treatment groups, most likely because neurons are the most highly sampled cell type in brain snRNA seq studies.

**Figure 2.**
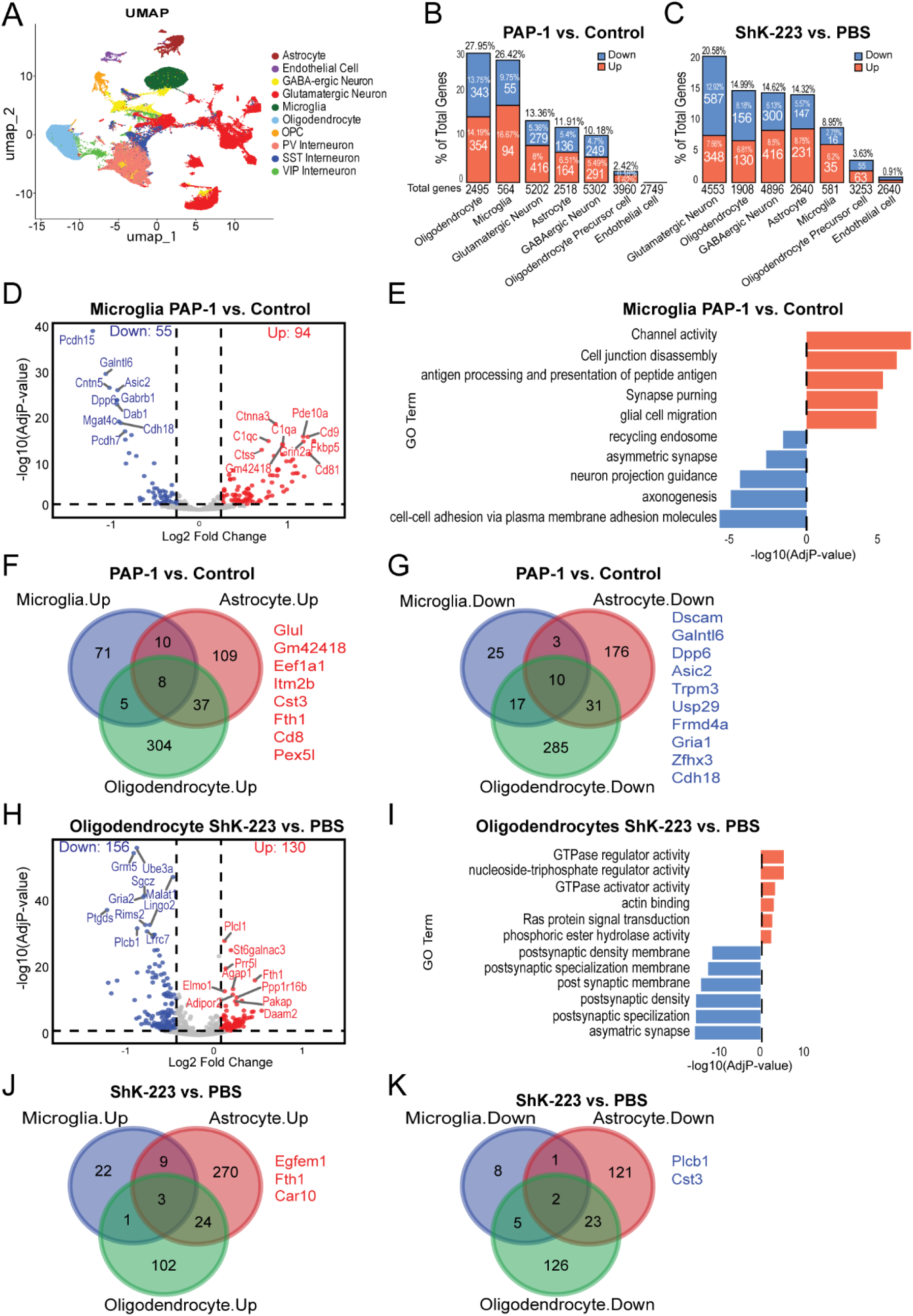
snRNA seq analysis of brain nuclei reveals distinct effects of PAP-1 and ShK-223 on microglia, astrocytes, and oligodendrocytes. **(A)** UMAP embedding of 23 transcriptionally distinct clusters derived from single-nucleus RNA-seq of 137,375 nuclei from (n= 3 per group, PAP-1, control chow, i.p ShK-223, i.p saline) and WT mice (n=2, no treatment). Each point corresponds to one nucleus, colored by its cell type identity. Cell types were assigned using marker genes from Sharma et al and Zhang et al.; each cluster annotated as Astrocytes (clusters 8, 20, 21), Microglia (6, 10), Endothelial cells (16), Oligodendrocytes (2), OPCs (22), GABAergic neurons (0, 5, 12, 13, 14, 18), and Glutamatergic neurons (1, 3, 4, 7, 9, 15, 17, 19, 23). **(B & C)** A comparison of proportions of differentially expressed genes (DEGs) per brain cell type: Stacked bar plots showing the proportion of upregulated (red) and downregulated (blue) DEGs across major brain cell classes in PAP-1 versus control **(B)** and ShK-223 versus PBS **(C)** experimental groups. Bars indicate the percentage of total significant genes per cell type, with numbers representing absolute gene counts. The y-axis represents the fraction of total DEGs relative to all genes detected per condition. **(D)** Microglial pseudobulk differential expression in PAP-1 versus control: Volcano plot showing significantly upregulated (red, *n* = 94) and downregulated (blue, *n* = 55) transcripts. The x-axis indicates log₂ fold change, and the y-axis represents –log^10^ adjusted *P* value. **(E)** GO enrichment of microglial DEGs in PAP-1 versus control: Bar plot showing significantly enriched Gene Ontology terms from pseudobulk microglial DEGs. The x-axis indicates –log^10^ adjusted *P* value; red bars represent pathways enriched among upregulated genes and blue bars among downregulated genes. **(F & G)** Overlap of DEGs across glial populations in PAP-1 versus control: Venn diagrams showing the intersection of upregulated (F) and downregulated (G) genes in microglia, astrocytes, and oligodendrocytes from pseudobulk snRNA-seq analysis. Shared and unique DEGs are indicated by circle overlaps. **(H & I)** Oligodendrocyte transcriptional and functional changes in ShK-223 versus PBS: **(H)** Volcano plot from pseudobulk snRNA-seq showing significantly upregulated (red, *n* = 130) and downregulated (blue, *n* = 156) oligodendrocyte transcripts. The x-axis represents log₂ fold change and the y-axis –log^10^ adjusted *P* value, with top DEGs labeled. **(I)** GO enrichment analysis of oligodendrocyte DEGs in Shk-223 treated 5xFAD mice. The x-axis indicates enrichment significance as -log^10^ adjusted P value; red bars correspond to pathways enriched among upregulated genes, while blue bars correspond to those enriched among downregulated genes. **(J & K)** Overlap of DEGs across glial populations in ShK-223 versus PBS: Venn diagrams showing shared and unique upregulated (**J**) and downregulated (**K**) genes in microglia, astrocytes, and oligodendrocytes from pseudobulk snRNA-seq analysis. Numbers indicate gene counts within each category, with representative DEGs listed for overlapping sets. See Supp. Fig. 2 for related supplemental analyses and figures.

We first focused on PAP-1 effects on glial populations (microglia, astrocytes, and oligodendrocytes). Pseudobulk analysis of microglial nuclei showed that PAP-1 treatment upregulated 108 and downregulated 65 DEGs **(Fig. 2D).** PAP-1 upregulated genes responsible for microglial effector functions (*CD81*), phagocytosis (*C1qa, C1qc, Cd9*), migration (*Robo2*) and plaque reduction (*Sorcs1*), while downregulated genes included *Cntn5, Asic2 and Usp29* which are known to induce adhesion and inflammatory pathways in microglia. Gene ontology (GO) enrichment of microglial DEGs also mirrored similar patterns with upregulated genes responsible for processes like cell junction disassembly, migration of cells and channel activity while downregulated genes include processes for cell-cell adhesion and axon genesis **(Fig. 2E)**. While PAP-1 did not systematically impact DAM or homeostatic gene signatures of microglia, PAP-1 treatment resulted in changes in microglial genes responsible for immunomodulatory, adhesion and migratory capabilities of microglia resulting in a protective phagocytic state of microglia which leads to better clearance of Aβ plaques. In astrocytes, PAP-1 treatment upregulated 273 genes involved in reducing astrogliosis and maintaining homeostasis (*Gpr37l1, Cst3, Ptgds*), Ab metabolism and clearance (*Clu* and *Apoe*). PAP-1 downregulated genes like *Kdm6a* and *lncGm26917*, known for their positive role in reactive astrogliosis and inflammation **(Supp. Fig. 2D)**. GO analysis of astrocyte DEGs showed that PAP-1 increased amyloid precursor protein metabolic process and cognition while decreasing actin filament organization and cell-cell adhesion (**Supp. Fig. 2E)**. These findings suggest that PAP-1 treatment alters the reactive state of astrocytes by reduced astrogliosis and induction of an anti-inflammatory state resulting in a shift from a reactive to homeostatic state. In oligodendrocytes, PAP-1 resulted in 369 upregulated and 347 downregulated genes. PAP-1 upregulated genes included regulators of oligodendrocyte plasticity and myelination (*Mal, Sgk1, Klf9, Plp and Pol3re*)^47, 48^ (**Supp. Fig. 2F)**. GO terms increased by PAP-1 included structural constituent of myelin sheath and oligodendrocyte differentiation, further supporting that PAP-1 overall enhances the functional state of oligodendrocytes contributing to improved myelination. (**Supp. Fig. 2G).** We next assessed the unique and shared gene expression changes induced by PAP-1 across these three glial subtypes **(Fig. 2F & 2G).** While the overlap among all three-cell types was relatively small (8 genes upregulated and 10 genes downregulated), PAP-1 related DEGs showed much greater overlap between astrocytes and oligodendrocytes (37 upregulated, 31 downregulated) indicating shared responses in these two glial cell types. Notably, the upregulated shared genes between the three glial cell types are responsible for iron metabolism *(Fth1*), migration (*CD81*), translation (*Eef1a1*), processing and Abeta clearance (*Cst3, Itm2B*) while shared downregulated genes are primarily responsible for cell adhesion (*Cdh18* and *Dscam*), anti-inflammatory functions (*Asic2* and *Usp29*) cell signaling (*Frmd4a*, *Trpm3*) and regulation of ion channel functions (*Dpp6*).

Next, we assessed the effects of ShK-223 treatment which resulted in the strongest transcriptional response in oligodendrocytes among the glial cells followed by astrocytes and microglia with 130 upregulated and 156 downregulated genes **(Fig. 2H).** The upregulated genes included several genes related to oligodendrocyte development and differentiation (*Ppp1r16b, Zbtb16*), cytoskeleton remodeling through GTPases (*Daam2, Elmo1and Agap1*) for formation of compact myelin sheath, RNA processing and transcription (*Polr3e and RbFox2*) while downregulated genes were critical regulators of proinflammation (*Ptgds*), synaptic spine structure (*Lrrc7*) and inhibitors of oligodendrocyte differentiation (*Prnp*). Interestingly, both PAP-1 and SK-223 showed improved functional state of oligodendrocytes by upregulation of oligodendrocyte differentiation and myelination related genes, though there was not much overlap in these upregulated genes. Upregulated GO terms also suggested similar functions like actin binding, GTPase regulator, and GTPase activator activity while downregulated terms include post synaptic membrane and asymmetric synapses **(Fig. 2I).** The effect of ShK-223 on microglia was modest, with 35 upregulated and 16 downregulated genes. Most of these upregulated genes include phagocytic (*Cd68*), adhesion (*Cdh8*) and noncoding RNA (*Bc1*) responsible for synaptic plasticity while downregulated genes include *Kctd12* and *Grik2* which are subunits of GABA-B and ionotropic glutamate receptors with known role in microglial activation (**Supp. Fig. 2H).** GO analysis mirrored similar patterns, with increased cell adhesion molecule binding and regulation of synapse organization, and decreased neurotransmitter receptor activity which suggest that ShK-223 treatment also resulted in a shift towards a protective microglial phenotype with changes in the overall functional capabilities of microglia. (**Supp. Fig. 2I).**

We also looked at the shared effects of both these inhibitors on microglia which showed again upregulation of iron metabolism (*Fth1*), cell motility, and survival (*Pak7*) (**Supp. Fig. 2J)**. Similarly, astrocytes showed increased expression of major NMDAR (*Grin2a*) and AMPA (*Gria3*) receptor subunits along with cytoskeleton rearrangement (*Elmo1*) but decreased expression of neuromodulator gene (*Ptgds*), vascular homeostasis (*Vegf, Ednrb*), and regulator of extracellular matrix (*Itih3*). (**Supp. Fig. 2K & 2L).** Further, we analyzed the overlap of the genes on ShK-223 treatment which showed that *Egfrem1*, *Fth1* and *Car10* were upregulated, while *Plcb* and *Cst3* were downregulated among glial cells **(Fig. 2J & 2K)**. Interestingly, we found *Fth1* to be a common upregulated gene upon treatment with either PAP-1 or ShK-223 in all glial cell types. Overall, our snRNA seq analysis of glial cells post Kv1.3 blockade using either PAP-1 and ShK-223 shows protective cell type specific effects with few shared genes showing up due to cellular and functional heterogeneity of these glial cells and different mechanisms of action of these blockers.

### Distinct effects of Kv1.3 blockade on excitatory and inhibitory neurons

We next investigated the effect of Kv1.3 blockers on transcriptional profiles of neurons, based on broad classification of neuronal nuclei into glutamatergic (excitatory) and GABAergic (inhibitory) neuronal classes. We analyzed the transcriptomic profile of glutamatergic neurons upon PAP-1 treatment which resulted in 416 upregulated and 279 downregulated genes. Notable upregulated genes are responsible for regulation of synaptic plasticity through cytoskeleton modulation (*Tpm1, Lrrk2*), formation of dendritic spine (*Homer1*), maintenance of membrane potential (*Atp1a1, Atp1b1 and Atp2b1*) and glutamate transportation (*Slc17a7*), while downregulated genes are responsible for regulation of neuronal differentiation (*Fgf1, Mtss1*) and gene regulation (*Hap1, Zep423, ZfhX3*) **(Fig. 3A).** GO analysis also mirrored similar patterns with upregulated terms like regulation of synapse structure or activity and dendritic spine **(Fig. 3B).** Interestingly, the transcriptional response to ShK-223 was stronger in glutamatergic neurons with 348 upregulated and 587 downregulated genes. Most of the highly upregulated genes included genes responsible for regulation of synaptic vesicles (*Sv2b*), synaptic plasticity (*Satb2*) and synaptic development (*Hs3st4*) suggesting a protective effect of ShK-223 on overall synaptic health. Notable downregulated genes are responsible for ubiquitination (*Ube3a*) and cytoskeleton regulation (*Bsg*) **Fig. 3G & 3H).** We also looked at the shared effects of both inhibitors on glutamatergic neurons which showed *Slc17a7* (glutamate transporter) to be concordantly upregulated while *Hap1* (vesicular transport) was concordantly downregulated **(Fig. 3F).**

**Figure 3.**
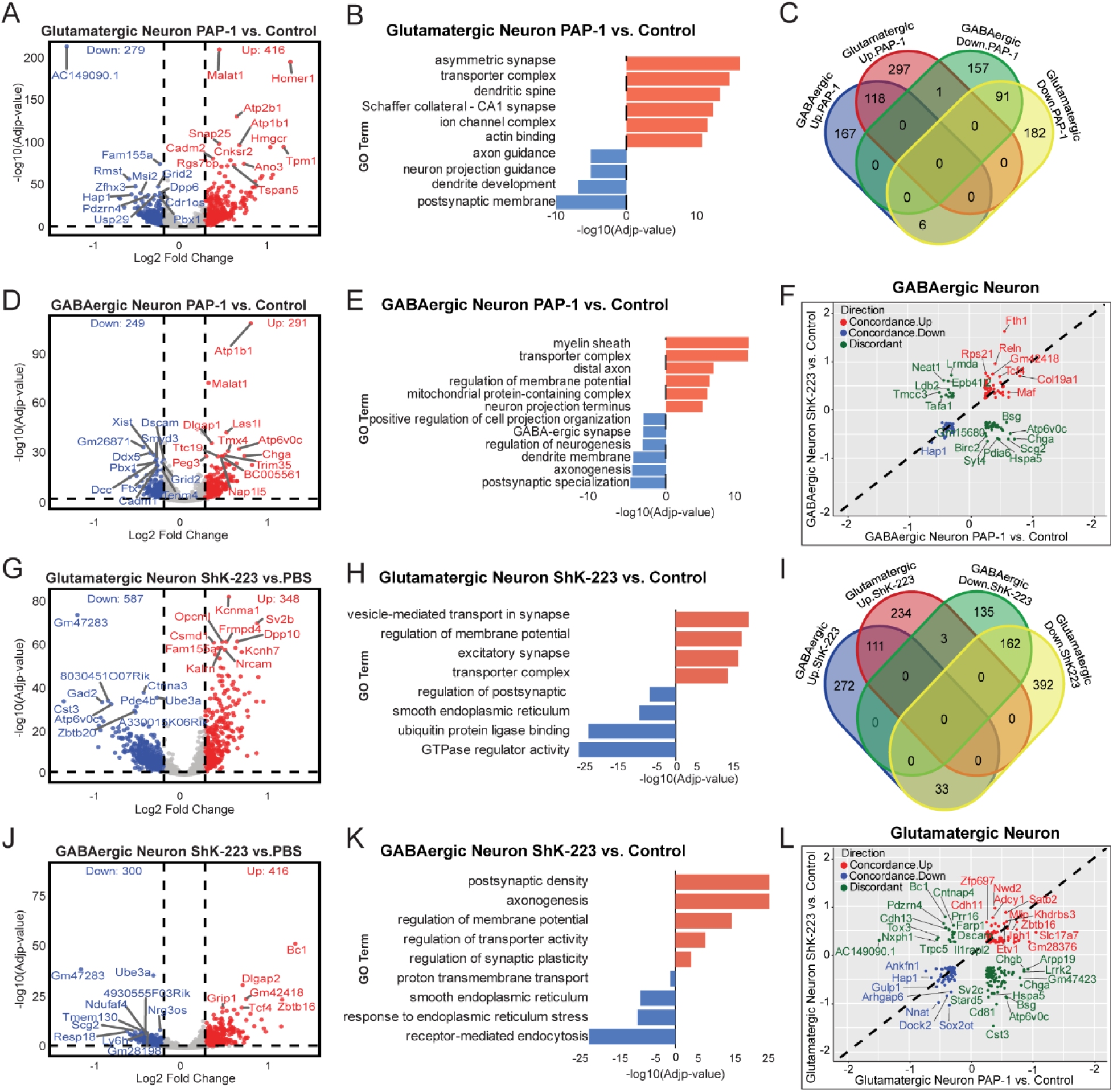
Differential effects of Kv1.3 blockade by PAP-1 and ShK-223 of glutamatergic and GABAergic neurons in 5xFAD brain. **(A)** Volcano plot representing DEGs comparing PAP-1 vs. Control in GABAergic neurons. DEGs with adjusted p-value < 0.05 and log₂FC > 0.25 are shown in red as upregulated, while DEGs with adjusted p-value < 0.05 and log₂FC < –0.25 are shown in blue as downregulated. **(B)** GO enrichment analysis of DEGs in GABAergic neurons, in PAP-1 vs. Control comparisons, shows significant pathways. In the bar plots, the X-axis represents _log10_(adjusted p-value) of the enriched pathways, while the Y-axis lists the pathway names. Each bar is sorted in descending order based on enrichment score. **(C)** Venn diagram showing shared and distinct upregulated and downregulated genes in GABAergic and glutamatergic neurons following PAP-1 treatment. **(D):** Volcano plot representing DEGs comparing PAP-1 vs. Control in Glutamatergic neurons. DEGs with adjusted p-value < 0.05 and log₂FC > 0.25 are shown in red as upregulated, while DEGs with adjusted p-value < 0.05 and log₂FC < -0.25 are shown in blue as downregulated. **(E)** GO enrichment analysis of Glutamatergic neurons in PAP-1 vs. Control shows significant enrichment pathways. In the bar plots, the X-axis represents log10(adjusted p-value) of the enriched pathways, while the Y-axis lists the pathway names. Each bar is sorted in descending order based on enrichment score. **(F)** Volcano plot showing differentially expressed genes in glutamatergic neurons following ShK-223 treatment versus PBS. **(G)** GO enrichment in glutamatergic neuron for ShK-223 (**H)** Venn diagram showing the overlap of upregulated and downregulated genes in GABAergic and glutamatergic neurons following ShK-223 treatment. **(I)** Volcano plot showing differentially expressed genes in GABAergic neurons following ShK-223 treatment versus PBS. **(J)** GO enrichment analysis of GABAergic neuron DEGs in ShK-223 vs PBS. See Supp Fig 3 for related supplemental analyses and figures.

Upon PAP-1 treatment, GABAergic neurons showed 291 genes significantly upregulated and 249 downregulated genes. Out of these DEGs, upregulated genes are known to be responsible for assembly and maturation of parvalbumin inhibitory synapse and interneurons (*Col19A1, Maf*), cytoskeleton remodeling (*Actg1, Bsg*) and regulation of membrane potential (*Atp1b1, Atp6voc*) while downregulated genes include genes regulating neurotransmitter release (*Baiap3*), axon guidance (*Dscam, Dcc*) and extracellular matrix (*Thsd4*) **(Fig. 3D).** GO analysis also revealed myelin sheath, mitochondrial protein containing complexes and regulation of membrane potential as upregulated terms **(Fig. 3E).** These changes indicate enhanced protective and restorative functions in the neurons. Similarly, ShK-223 treatment also resulted in a robust transcriptional response with several upregulated genes responsible for improved synaptic function (*Reln, Slc1a2*) and neuronal polarization and migration (*Fgf13*), while downregulated genes are responsible for cellular protein homeostasis (*Ube3a*) and quality control within endoplasmic reticulum (*Hspa5*) **(Fig. 3J).** GO analysis also showed regulation of synaptic plasticity and axon genesis as upregulated while downregulated terms included response to endoplasmic reticulum stress **(Fig. 3K).** Notably, ShK-223 also highly upregulated some genes in common with PAP-1, including *Col9a1* and *Fth1* **(Fig. 3L).** Further, we subclassified glutamatergic and GABAergic neurons into subclusters (8 Glutamatergic neuron subclusters: 1, 3, 4, 7, 9, 15, 17, and 19; and 6 GABAergic subclusters: 0, 5, 12,13, 14, and 18) and examined the effects of PAP-1 and ShK-223 on proportions of neuronal subclusters. We found that both drugs altered proportions of neuronal subclusters. Specifically, both drugs increased proportions of excitatory neuronal cluster 1. **(Supp. Fig. 3E & 3F)**

Next, we assessed the overlap in the effects of Kv1.3 blockade on excitatory and inhibitory neurons. In the case of PAP-1 treatment, we found 118 upregulated genes common between glutamatergic and GABAergic neurons in comparison to 91 downregulated genes common genes, which is a significant overlap between the two subtypes **(Fig. 3C).** GO analysis also showed common upregulated terms like transporter complex and downregulated terms like postsynaptic membrane/specialization. Conversely, ShK-223 treatment resulted in more downregulated (162 genes) compared to upregulated (114 genes) common between the two neuronal subtypes with common GO terms like transporter complex (upregulated) and smooth endoplasmic reticulum (downregulated) **(Fig. 3H)**. The percentage of common genes differentially expressed between the two subtypes is half or one third in some cases compared to the total DEGs which means these blockers have significant conserved effects on both neuronal classes involving synaptic health, energy metabolism, and plasticity, although class-specific effects are also present. Overall, these neuron-specific effects of PAP-1 and ShK-223 are consistent with protective effects of Kv1.3 blockers on synaptic health, intracellular homeostasis, neuronal excitability, and neurotransmission.

### Cell-cell communication analysis reveals augmentation of neuro-vascular coupling by Kv1.3 blockade

Single nuclear RNA seq data allows an assessment of ligand-receptor pairs expressed by different cell types, to obtain insights into intercellular communication among neuronal, glial, and vascular cell types. Based on the number of ligand-receptor pairs (L/R pairs) expressed across cell type pairs, a probability of interaction can be estimated. We established cellular networks using pairs of membrane receptors and their ligands to gain molecular insights into this cross cellular signaling upon either PAP-1 or ShK-223 treatment. We found that PAP-1 treatment increased communication probability between glutamatergic neurons and GABAergic neurons with endothelial cells, suggesting increased cellular crosstalk between neuronal and vascular elements **(Fig. 4A)**. Neuro-vascular L/R pairs most upregulated by PAP-1 included SLIT2/ROBO signaling, where Slit2 was increased in endothelial cells while out of its receptors present on GABAergic neurons (Robo1/Robo2) and Glutamatergic neurons (Robo2), only Robo2 was upregulated in GABAergic neurons^49^. This signaling axis results in migration and polarization of endothelial cells, suggesting the impact of PAP-1 on neurovascular coupling. In contrast, PAP-1 reduced signaling between PTN and its receptors PTPRZ1 and ALK on GABAergic neurons **(Fig. 4B).** ShK-223 treatment resulted in increased communication probability between astrocytes and glutamatergic and GABAergic neurons **(Fig. 4C).** Specifically, ShK-223 increased signaling between astrocytic Nrxn1 and ligands (Nlgn1, Lrrtm4, Lrrtm3, Clstn2) in both Glutamatergic and GABAergic neurons (**Fig. 4D**). Both neuroligin (NLGN1) and neurexin (NRXN1) are expressed by astrocytes and neurons, and their bidirectional signaling is known to play a role in synaptogenesis and astrocytic morphogenesis^50^. Interestingly, NRXN1 (pre-synaptic) and LRRTM4 (post-synaptic) interaction was increased in GABAergic neurons, but were downregulated in glutamatergic neurons, again suggesting distinct roles of this pair in synaptogenesis in two classes of neurons **(Fig. 4G & 4H).**

**Figure 4.**
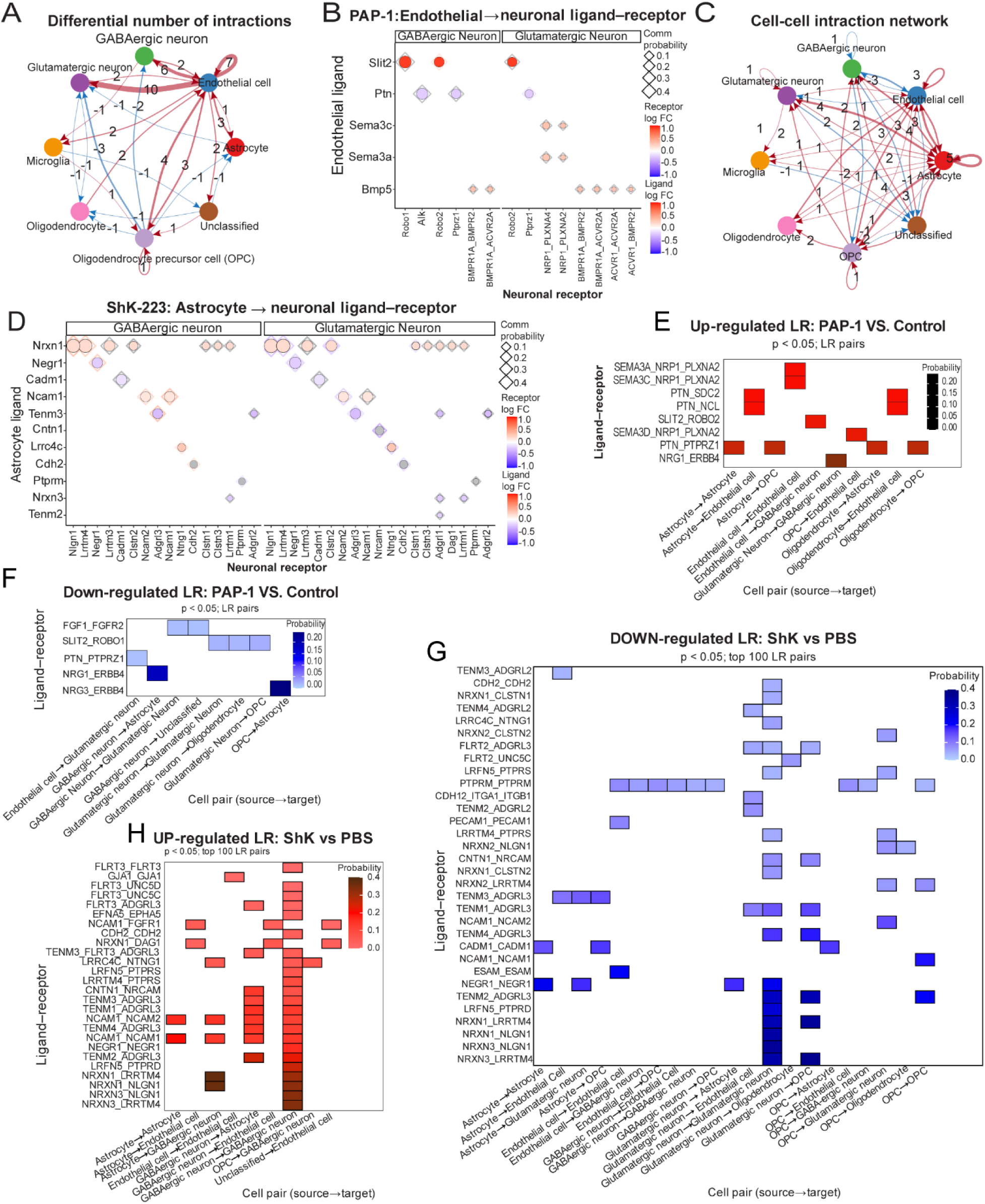
Cell-cell communication networks are altered by Kv1.3 Blockade. **(A)** Cell-cell communication network after PAP-1 treatment: Nodes represent cell types, and arrows indicate the direction of ligand receptor signaling, with thicker lines reflecting stronger interactions. PAP-1 increases communication mainly among glial populations, including astrocytes, microglia, oligodendrocytes, and OPCs, with additional signals directed from glutamatergic and GABAergic neurons toward astrocytes and endothelial cells. Line colors correspond to specific signaling routes between cell populations, helping distinguish outgoing versus incoming interactions visually. Numbers displayed on the connections indicate the total count of ligand receptor pairs identified between those two cell types. **(B)** Endothelial to neuronal ligand receptor interactions following PAP-1 treatment are shown for GABAergic and Glutamatergic neurons. The y-axis denotes endothelial ligands, and the x-axis denotes neuronal receptors. Diamonds shape represents receptor log fold change (log FC), while circles represent ligand log FC. Symbol size corresponds to communication probability, and color indicates the direction and magnitude of log FC**. (C)** Compared to PAP-1, ShK-223 produces a more extensive and interconnected network, with strong bidirectional signaling among glutamatergic and GABAergic neurons, oligodendrocytes and OPCs, and multiple pathways linking neurons with glial cells. This pattern reflects a broader remodeling of cell–cell communication. **(D)** Astrocyte to neuronal ligand receptor interactions following ShK-223 treatment is shown for GABAergic and Glutamatergic neurons.. The y-axis denotes endothelial ligands, and the x-axis denotes neuronal receptors. Diamonds shape represents receptor log fold change (log FC), while circles represent ligand log FC. Symbol size corresponds to communication probability, and color indicates the direction and magnitude of log FC. **(E)** Heatmap based on CellChat analysis displaying up-regulated ligand-receptor pairings (p < 0.05) in PAP-1 vs. Control. A significant interaction between cell types that express ligands (source) and receptors (targets) is indicated by each rectangle; the probability of communication is reflected in the color of intensity. **(F)** Heatmap of PAP-1 vs. Control down-regulated ligand-receptor pairs (p < 0.05), where a blue gradient denotes a lower likelihood of communication. **(G)** Heatmap of up-regulated ligand-receptor pairs (p < 0.05) in ShK-223 vs PBS, with red gradient denoting higher communication probability. **(H)** Heatmap of down-regulated ligand-receptor pairs (p < 0.05) in ShK-223 vs PBS, with blue gradient indicating reduced probability of ligand-receptor signaling.

We also identified several L/R pairs that were concordantly up- or down-regulated, representing higher confidence pathways of cell-cell communication that were altered by PAP-1 (**Fig. 4E & 4F**) or ShK-223 (**Fig. 4G & 4H**) treatment. The L/R pair NRG1-ERBB4 showed the highest upregulation in glutamatergic and GABAergic neurons, where NRG1 ligand is predominantly expressed by glutamatergic neuron and its receptor ERBB4 is expressed on GABAergic neurons. Previous studies have shown that excitatory–inhibitory (*E*–*I*) balance in cortex is regulated by NRG1-ERBB4 L/R pair^51^. Hence, the upregulation of NRG1-ERBB4 suggests that PAP-1 treatment increases this signaling axis to restore the known (E-I) imbalance in 5xFAD mice^52^. In contrast to increased signaling in glutamatergic and GABAergic neurons, NRG1-ERBB4 showed decreased signaling in astrocyte-GABAergic neuron communication, suggesting a cell type specific role of this interaction in maintenance of glutamate and GABA balance. Further, the PTN-PTPRZ1 L/R pair involved in Wnt/β-catenin axis was found to be upregulated in non-neuronal cell type combinations including astrocyte-oligodendrocyte, astrocyte-OPC, astrocyte-astrocyte and oligodendrocyte-OPC in PAP1 treated mice. PTN-PTPRZ1 has been shown to play an important role in remyelination, oligodendrocyte survival, and astrocyte morphogenesis^53^. This further confirms our pseudobulk analysis findings that PAP-1 treatment leads to upregulation of myelination by oligodendrocytes along with changes in functional state of astrocytes, and that the PTN-PTPRZ1 Wnt/β catenin signaling axis may play a role in these changes. Overall, both PAP-1 and ShK-223 treatment results in activation of different L/R pairs among different cell types which can be the basis of profound effect of these drugs on various brain cell types ultimately resulting in their cumulative protective effects.

### Bulk brain transcriptomics and proteomics with PAP-1 confirm increased levels of synaptic proteins and decreased glial activation

Since our single nuclei analysis showed a profound effect of PAP-1 treatment on all the major glial cell types and neurons in 5xFAD mice we next assessed changes on the bulk brain transcriptome and proteome. Bulk-omics also complement snRNA seq data by deeper coverage of the transcriptome, allowing more robust upstream pathway analyses. Global transcriptome analysis of control and PAP-1 treated 5xFAD mice, after removal of low-abundance transcripts, resulted in 14,662 mRNAs. Differential expression analysis of these genes was performed using DESeq2, and significance was determined using Benjamini-Hochberg adjusted p-values which revealed that PAP-1 treatment significantly induces both upregulation (212 genes) and downregulation (175 genes) of several genes **(Fig. 5A).** Among the top upregulated genes are *C1ql2, Slc9a4, Bdnf, Fosl2, Arc, Nlgn1* and *Nptxr* which are known to be involved in processes such as synaptic signaling, membrane trafficking, and neuronal plasticity^54, 55, 56^. GO analyses of DEGs showed enrichment of terms like synapse assembly, synaptic vesicle membrane, membrane adhesion, and retrograde trans-synaptic signaling, which suggests a transcriptional shift to promote synaptic organization, neurotransmission, and neuronal communication by PAP-1 **(Fig. 5B).** In contrast, the top downregulated genes included *Nexn, Adamts, Lrrc1, Srgap* and *Lingo3* which are involved in glial regulation, extracellular matrix degradation along with cytoskeleton remodeling and show enrichment of pathways related to gliogenesis, glial cell differentiation, Schwann cell differentiation in GO analysis. We also performed Ingenuity Pathway Analysis (IPA) to nominate the differentially regulated pathways responsible for these observed transcriptomic changes **(Fig. 5C).** Our analysis resulted in similar upregulated signaling like synaptogenesis, neurovascular coupling, and GABA receptor activation along with specific pathways like Netrin-1 signaling, Wnt/β NGF stimulated transcription and GABA receptor activation with known role in these cellular changes^57, 58^. The downregulated pathways included IL3, JAK/STAT, NGF, and CREB signaling further suggesting the modulation of cellular and immune functions of the different brain cells. Together, these results point to increased synaptic remodeling and neurotransmission, and decrease in gliogenesis and associated immune pathways, because of PAP-1 treatment. These overall synaptic effects of PAP-1 observed in the bulk transcriptome, were consistent with snRNA seq pseudobulk analyses of excitatory and inhibitory neurons.

**Figure 5.**
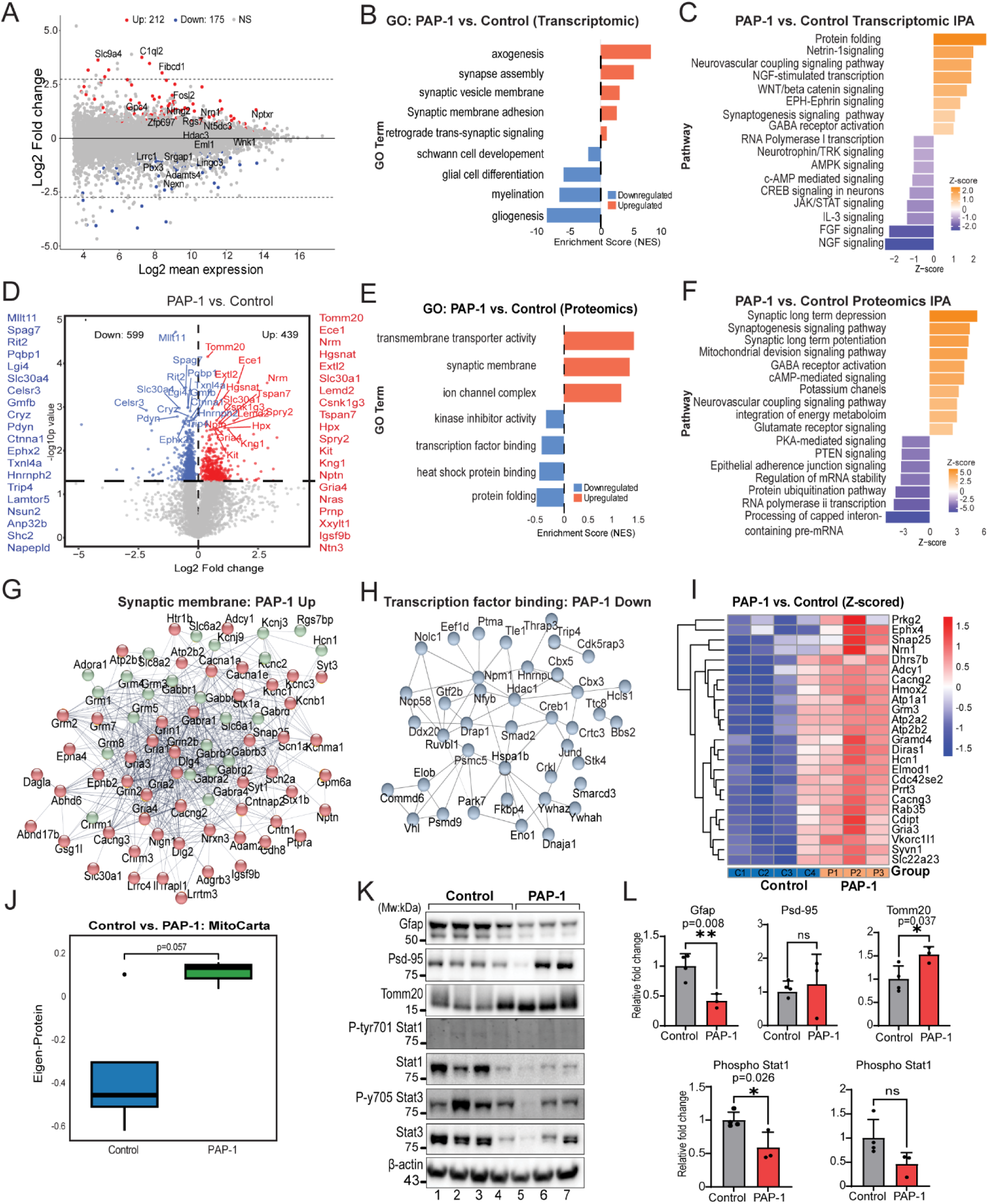
Effects of Kv1.3 blockade on brain transcriptomes and proteomes. **(A)** MA plot showing differential gene expression between PAP-1-treated and control samples. The y-axis displays log₂ fold change (Log₂ FC), and the x-axis shows log₂ mean expression for each gene. Each dot represents a gene: red indicates significantly upregulated (n = 212), blue indicates significantly downregulated (n = 175), and yellow denotes genes with no significant change (NS) at adjusted p ≤ 0.05. Horizontal dashed lines mark the fold-change thresholds (±2.5 Log₂ FC). (**B)** GO enrichment analysis of bulk transcriptomic changes in PAP-1 versus Control. The x-axis shows normalized enrichment score (-log^10^ adjusted p-value ≤ 0.05), and the y-axis lists enriched pathways. Bar length reflects enrichment strength, with pathways ranked in descending order with upregulated categories in red whereas downregulated categories in blue (**C)** IPA of transcriptomic changes in PAP-1 versus Control. The x-axis represents Z-scores, and the y-axis lists enriched pathways. Orange bars indicate predicted activation (positive Z-scores), while blue bars indicate predicted inhibition (negative Z-scores). Bar length corresponds to the strength of predicted activity. (**D)** Volcano plots of proteomic changes between PAP-1 and Control identified using one-way ANOVA. The x-axis shows log₂ fold change, and the y-axis shows -log^10^ P-value. Significantly upregulated proteins (n = 439) are shown in red, significantly downregulated proteins (n = 599) are shown in blue, and non-significant proteins are shown in grey (FDR-adjusted p ≤ 0.05). Representative significant hits are labeled. **(E)** GO analysis of proteomic differences between PAP-1 and Control. The x-axis represents normalized enrichment score (NES, FDR-adjusted p ≤ 0.05), while the y-axis lists the enriched biological terms. **(F)** IPA of proteomic changes in PAP-1 versus Control. The x-axis represents Z-scores, and the y-axis lists significantly enriched pathways. **(G)** Protein-protein interaction network of synaptic membrane proteins enriched in PAP-1. Nodes represent proteins, and edges indicate interactions. The cluster shows strong connectivity among synaptic receptors, ion channels, and scaffolding proteins, with central nodes including GRIN1, GRIN2A, GABRA1, SNAP25, and CACNA1A. **(H)** Protein-protein interaction network of transcription factor binding proteins enriched in PAP1. The cluster centers on regulatory hubs such as NPM1, CREB1, HDAC1, and HSP1A1, reflect coordination among transcriptional regulators. **(I)** Heatmap showing Z-scored expression of resilience-associated proteins, highlighting consistent upregulation in PAP-1–treated samples compared with controls. **(J)** Box plot showing enrichment of mitochondrial proteins (MitoCarta set) in Control versus PAP-1. The y-axis represents an enrichment score. PAP-1 samples display a significant increase compared to control, indicating enhanced mitochondrial protein abundance. (**K & L**) Validation of protein abundance using western blot and densitometric analysis of markers of synaptic integrity, gliosis and neuroinflammation along with mitochondrial protein TOMM20. See Supp. Fig. 4 for related supplemental analyses and figures.

Next, we assessed the global proteomic changes in the 5xFAD mouse brain post-treatment and assessed differential abundance of proteins between the control and PAP-1 treated group. Of 6,670 quantified proteins, 439 proteins were increased while 599 proteins were decreased by PAP-1 treatment **(Fig. 5D).** The top upregulated proteins included TOMM20 (mitochondrial transporter), ECE1 (amyloid beta degradation), NRM (neuronal homeostasis), NPY1R (neuropeptide receptor) and SLC30A1 (Zinc transporter and neuroprotection), while downregulated proteins included MLLT11(chromatin reader for efficient RNA polII transcription), PQBP11(splicing and transcription) and RIT2, ATG12 and ATG14 (autophagy). GO analyses also showed upregulation of terms like transporter activity and synaptic membrane (75 proteins), while downregulated terms included transcription factor binding (40 proteins) and protein folding including stress response **(Fig. 5E).** We also identified protein-protein interactors (STRING analysis) within 75 synaptic membrane proteins increased with PAP-1, and 40 Transcription binding proteins decreased by PAP-1 **(Fig. 5G & 5H).** In the protein-protein interactions of synaptic membrane, we found two functionally distinct subsets, namely, intrinsic component of synaptic membrane (52 proteins) like glutamatergic proteins involving receptors, subunits GRIA2, GRIA3 and regulation of postsynaptic membrane potential (23 proteins); as well as a cluster of GABAergic proteins like GABBR1 and GABRB2. Protein-protein interactions of transcription factors binding proteins downregulated by PAP-1 included SMAD2 and JUND, which play a key role in cellular differentiation, proliferation, and immune signaling. Further, pathway analysis of bulk proteomes from PAP-1 treated mice identified mitochondrial division, integration of energy metabolism and neurovascular coupling as increased by PAP-1, while RNA pol II transcription and regulation of mRNA stability were decreased by PAP-1 **(Fig. 5F).** Strikingly, neurovascular coupling was also predicted by cell-cell communication analysis of snRNA seq data, further highlighting the importance of this pathway as a mechanism of neuroprotection by Kv1.3 blockers. The IPA analysis of both bulk transcriptome and proteome from PAP-1 and control treatment groups, also identified common pathways such as synaptogenesis and GABA receptor activation, which also suggests the protective effect of PAP-1 on synapse restoration and neuronal communication. However, there are several differences also highlighted in the IPA analysis of transcriptome and proteome that can further be explained by modest correlation between the treatment effects observed at the level of the bulk transcriptome and proteome **(Supp. Fig. 4A)**.

To further understand the cell type-specific upstream regulation of all these predicted biological consequences, we applied upstream regulator analysis in IPA to DEGs derived from Pseudobulk analyses of six major cell types in our snRNA seq based pseudobulk analysis. We identified several key upstream regulators in these cell types which are known to drive changes in the gene expression or modulate the function of other genes by modifying the cellular environment. In microglia, the upstream regulators of activated pathways include IFNAR, STAT5A/B, CDC42, MAP3K8, and C5AR1, which are responsible for increased immune surveillance, complement activation, and cell migration. Upstream regulators of downregulated pathways include GLB1, NEU3, PSEN1, and BACE1 responsible for amyloid β precursor processing and neuroinflammation **(Supp. Fig. 4C)**. Similarly, in oligodendrocytes TCF7L2, INSR, GPER1 are activated upstream regulators with their known role in myelination while APH1, which is downregulated, is responsible for processing of amyloid β precursor protein. **(Supp. Fig. 4E)**. These findings based on upstream regulators in glial cells also suggest consistent upregulation of myelination-related pathways while amyloid β processing is downregulated, resulting in lesser amyloid β production. Next, we assessed the upstream regulators in glutamatergic neurons which showed upregulation of factors like IL5 responsible for inhibition of proinflammatory genes and CD38 responsible for synaptogenesis by increasing extracellular levels of SPARCL1 **(Supp. Fig. 4G)**. Interestingly, in GABAergic neurons upregulation of HFE as an upstream regulator suggests a protective response to limit exposure of iron **(Supp. Fig. 4H)**. We also observed overlaps in several upregulated (NRF1, TEAD, NFE2L2) and downregulated (ZHX2, KDM5A) upstream regulators between two neuronal subtypes as expected due to large overlap in their shared DEGs.

Based on our findings of increased levels of synaptic and neurotransmission related proteins following Kv1.3 blockade in the brain, we hypothesized that Kv1.3 blockade augments mechanisms of cognitive resilience. Proteins that determine cognitive resilience (pro-resilience) have been previously identified by correlating post-mortem brain proteomes with rate of cognitive decline in a large longitudinal study of aging (ROSMAP)^59^. In this study, over 250 pro-resilience proteins were identified that are positively correlated with cognitive slope, along with over 300 anti-resilience proteins negatively associated with cognitive slope. We intersected pro-resilience and anti-resilience proteins lists with proteins that showed differential abundance due to PAP-1 treatment in bulk brain proteomics data **(Supp. Fig. 4I)**. We observed a more disproportionate enrichment of pro-resilience proteins in the group of proteins upregulated by PAP-1 than expected (Chi square 15.79, p=0.0004), suggesting a preferential effect of PAP-1 on increasing these resilience pathways in mouse brain (**Supp. Fig. 4J)**. 25 pro-resilience proteins were increased by PAP-1 in the brain proteome including NRN1, the top pro-resilience protein identified in humans **(Fig. 5I & Supp. Fig. 4K).** Interestingly, the expression levels of NRN1 also showed a significant increase in our bulk transcriptomic and pseudobulk analysis of glutamatergic neurons in the PAP-1 treated group **(Supp. Fig. 4L, 4M, 4N, 4O)**. These pro-resilience proteins increased by PAP-1 are shown in **Fig 5I** and include synaptic proteins (SNAP25, NRN1, PRKG2) and mitochondrial proteins (HMOX2). Based on these results, we infer that pro-resilience pathways are also contributing to the neuroprotective effect of Kv1.3 blockade.

Bulk proteome analysis of PAP-1 treatment effects identified several top differentially regulated mitochondrial proteins like TOMM20 and NRM. The proteome-based IPA also nominated the mitochondrial division along with integration of energy metabolism as an upregulated pathway suggesting the involvement of mitochondria. The presence of Kv1.3 in mitochondria is established and PAP-1 is a cell permeable Kv1.3 inhibitor, hence, we next interrogated the organelle specific effect of PAP-1 treatment on the expression of mitochondrial proteins listed from MitoCarta (Ver 3.0). We estimated an eigen protein to reflect mitochondrial proteins in each sample and found that mitochondrial proteins were generally increased in the PAP-1 treated group compared to control **(Fig. 5J)**. We further verified the increased levels of top upregulated mitochondrial protein TOMM20 by protein abundance in PAP-1 treated bulk proteome MS data and by western blot. **(Fig. 5K & 5L) (Supp. Fig. 4Q)**. Unlike PAP-1, ShK-223 treatment which upregulated 240 DEPs and downregulated 252 DEPs **(Supp. Fig. 4B)**, did not increase the mitochondrial proteins when compared to PBS-treated control, consistent with a preferential effect of PAP-1 treatment on increasing brain mitochondria protein abundance**.(Supp. Fig. 4P)** In bulk brain proteomics data, regulators of mitochondrial biogenesis (AMPK, mTOR), and fission and fusion were not impacted, while OPA3 (log2FC 0.73, p=0.029), a protein involved in mitochondrial maintenance, was increased by PAP-1.

To gain a further comprehensive understanding of different neuroprotective mechanisms exhibited by PAP-1, we looked at the abundance of protein markers of gliosis (GFAP) and synaptic integrity (SV2A and PSD-95) **(Supp. Fig. 4Q).** Further, we validated the expression levels of these proteins along with neuroinflammation driving transcription factors STAT1/3 and their phospho-forms by western blot **(Fig. 5K&5L) (Supp. Fig. 4R).** Protein abundances by MS and western blots showed decreased levels of markers of gliosis, and an increase in synaptic markers due to PAP-1. PAP-1 also decreased neuroinflammatory pathways as depicted by decreased levels of both phospho-and total STAT1/3 by western blot. These comprehensive bulk transcriptomic and proteomic findings provide evidence for protective effects of PAP-1 on neurons and synapses with a reduction in reactive astrogliosis and neuroinflammation, as indicated by decreased activation of STAT1/3.

### Kv1.3 interacts with STAT proteins and specifically regulates type 2 IFN signaling

In our previous work, we showed that Kv1.3 is highly co-expressed with pro-inflammatory IFN-related genes in microglia in AD mouse models^19^. In more recent work, we published a Kv1.3 interactome from BV2 microglial cells using proximity-labeling (by fusing TurboID to Kv1.3). We found that Kv1.3 lies in proximity with IFN-related proteins including STAT1/3^27^, which suggests Kv1.3 and interferon signaling may be functionally related. Further in **Fig. 5K**, we found inhibition of STAT1/3 phosphorylation in PAP-1 treated mouse brain lysate, supporting functional coupling between Kv1.3 and interferon signaling. Hence, we wanted to gain further mechanistic insights into the functional coupling of this ion channel with interferon signaling **(Supp. Fig. 5A)** since this has not been investigated in microglia. To first validate Kv1.3-STAT1/3 interactions in BV2 microglia cells, we leveraged V5 tag in our V5-TurboID-Kv1.3 fusion construct and performed V5 tag antibody-based co-immunoprecipitation followed by STAT1 and STAT3 immunoblots. Both STAT1 and STAT3 were found to co-immmunoprecipitate with Kv1.3 confirming a protein-protein interaction between STAT1/3 with Kv1.3 (**Fig. 6A).** To determine whether Kv1.3 channels regulate STAT1 and 3 signaling in response to interferons, we investigated the effects of Kv1.3 blockade by either PAP-1 or ShK-223 on microglial cells overexpressing Kv1.3, followed by IFN-γ (type 2) or IFN-α/β (type 1) induction. As expected, IFN-γ induction led to phosphorylation of STAT1 Tyr701, STAT3 Tyr 705 and JAK1 Tyr 1034. Blockade by PAP-1 prior to IFN-γ induction resulted in reduced phosphorylation of STAT1, STAT3 and JAK1 on these phosphorylation sites, with not much changes in the total proteins. PAP-1 mediated blockade of STAT1 phosphorylation at Tyr701 upon IFN-γ induction has also been reported earlier by our group^27^. Interestingly, the ShK-223 blockade had no significant effect on phosphorylation of these proteins **(Fig. 6B & 6C)**. Since, BV2 SR73 is a Kv1.3 overexpression system we wanted to check the functional coupling between Kv1.3 and IFN signaling in the MMC microglial cell line with endogenous Kv1.3 expression^21^. Here also only PAP-1 mediated blockade led to inhibition of phosphorylation of STATs (**Fig. 6D & 6E**). Since Kv1.3 blockade led to inhibition of IFN-γ induced phosphorylation, we next wanted to study the effects of Kv1.3 blockade with IFN-α induction. Surprisingly, with IFN-α induction, PAP-1 and ShK-223 blockade did not impact phosphorylation of STAT1, STAT3 and JAK1 in both cell types. (**Fig. 6F & 6G) (Supp. Fig. 5B & 5C).** As PAP-1 mediated pharmacological blockade of Kv1.3 showed an inhibitory effect on IFN-γ, we next assessed the effect of genetic deletion of Kv1.3 upon IFN-γ induction in primary postnatal microglia isolated from wild-type and *Kv1.3* KO pups. Consistent with our findings in microglial cell lines, we observed a significant reduction of STAT1 and STAT3 phosphorylation in response to IFN-γ in *Kv1.3*-KO microglia, as compared to wild-type microglia (**Fig. 6H & 6I**). Based on these *in vitro* studies using pharmacological and genetic approaches in cell lines and primary microglia, we conclude that Kv1.3 channels interact with STAT1 and STAT3, with specificity for type 2 IFN but not type 1 IFN signaling.

**Figure 6.**
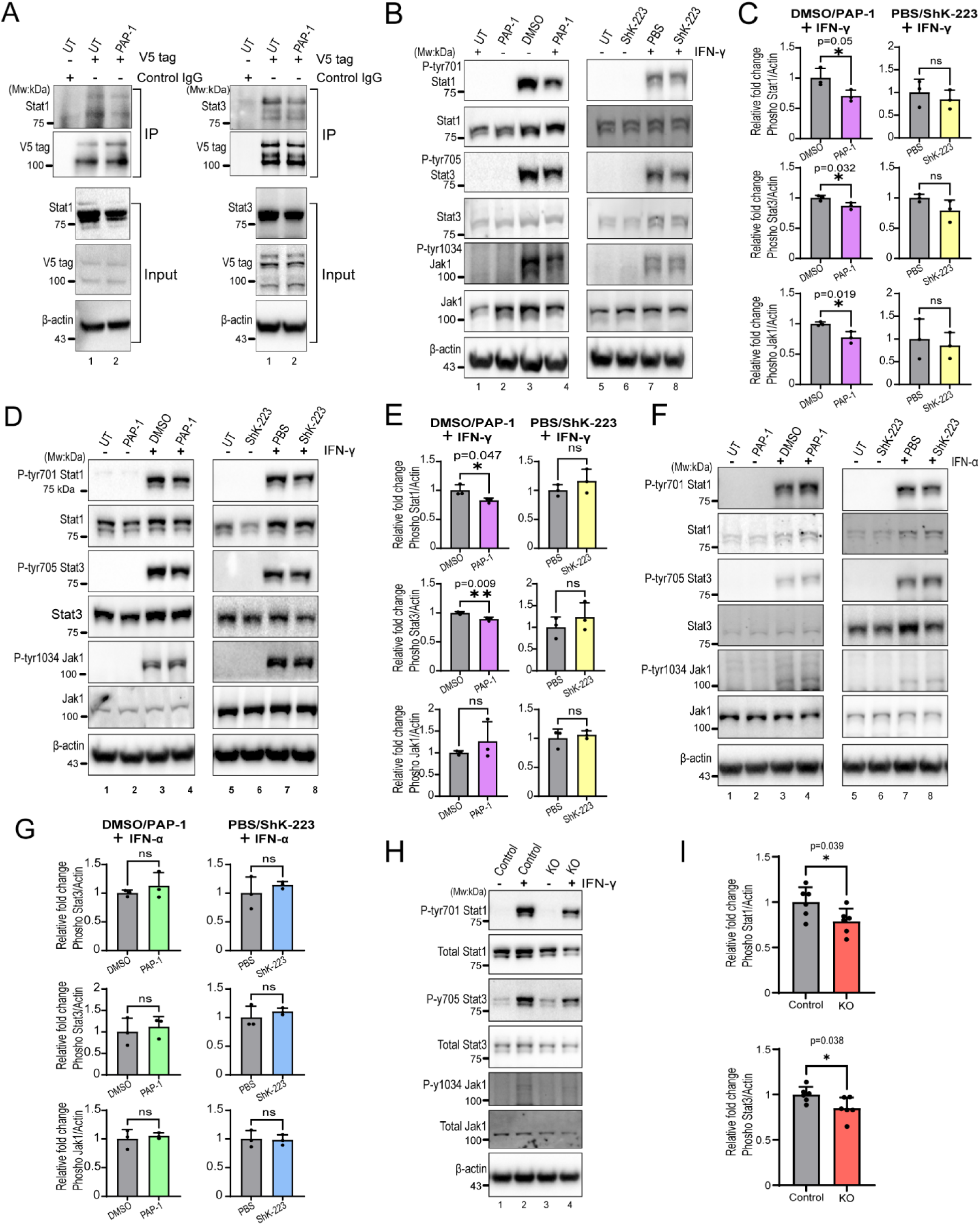
Kv1.3 inhibition by PAP-1 specifically downregulates type 2 IFN signaling. **(A)** Coimmunoprecipitation followed by western blot of whole cell lysate of BV2 cells confirmed the physical interaction between V5 tagged Kv1.3 and STAT1/STAT3. **(B & C)** Western blot and densitometric analysis of whole cell lysate of interferon γ induced BV2 cells post Kv1.3 blockade by PAP-1 shows reduced phosphorylation of STAT1, STAT3 and JAK11. ShK-223 blockade showed visual reduction but not statistically significant. **(D & E)** Western blot and densitometric analysis of whole cell lysate of interferon γ induced MMC cells post Kv1.3 blockade by PAP-1 shows reduced phosphorylation of STAT1 and STAT3 with no changes in JAK11. ShK-223 blockade showed no effect. **(F & G)** Western blot and densitometric analysis of whole cell lysate of IFN-α induced BV2 cells post Kv1.3 blockade by PAP-1 or ShK-223 shows no effect on phosphorylation of STAT1, STAT3 and JAK1. **(H & I)** Western blot and densitometric analysis of whole cell lysate of IFN-γ induced primary microglia cells from WT vs *Kcna3* KO mice post Kv1.3 blockade by PAP-1 shows reduced phosphorylation of STAT1 and STAT3. See Supp. Fig. 5 for related supplemental analyses and figures.

### Brain pathological effects of Kv1.3 blockade by PAP-1 and ShK-223 are reflected in cerebrospinal via AD-relevant biomarkers of therapeutic effect

CSF is considered a window into brain pathological changes. Disease effects occurring in the brain are often reflected in CSF, allowing less-invasive monitoring of brain pathological changes in response to disease or disease-modification using CSF biomarkers. Since prior work and this study confirm disease-modification by Kv1.3 blockers based on decreased plaque pathology and neuroprotective and restorative molecular changes, we performed DIA MS on CSF collected from experimental animals at time of euthanasia. DIA MS of CSF identified 2,495 proteins, 2,227 proteins of which were also identified in brain proteomes while 268 proteins were unique to the CSF. These 268 proteins were enriched in ECM, immune/blood/coagulation/complement/endopeptidase related terms. After removing the effect of blood contamination, we compared CSF proteomes across experimental groups **(Supp. Fig. 6A-F).** 429 proteins showed pathology-associated changes (283 increased and 146 decreased in 5xFAD mice compared to WT controls **(Fig. 7A).** Proteins increased in CSF involved lysosomal and immune/myeloid related mechanisms, consistent with increased immune activation and lysosomal dysfunction in the brain as a result of progressive amyloid pathology **(Fig. 7D).** As compared to the control group, PAP-1 treated mice showed increased levels of 28 proteins (including ATP6V1E1, SMS and CZIB) and decreased levels of 11 proteins (including VAT1L, NEFL and NDRG1) **(Fig. 7B).** Level of NEFL, a biomarker that reflects axonal damage across neurological disorders^60^, was reduced by PAP-1, consistent with disease modification observed in the brain. GO analysis showed the upregulation of transporter activity and SH3 binding domain while downregulation of endosome membrane as effects of PAP1 treatment in CSF **(Fig. 7E).** ShK-223 upregulated 46 proteins (including LASP1, PROC and PREPL) and downregulated 137 proteins (including ISYNA1, UBA7 and ISG15) in CSF **(Fig. 7C).** Of these, ISG15, a protein that reflects interferon-mediated responses, was decreased by ShK-223 treatment. GO terms increased in the CSF by ShK-223 included regulation of homeostasis and migration of endothelial cells while those decreased in CSF included glycolytic process and fatty acid binding **(Fig. 7F).** Interestingly, we observed minimal overlap between CSF DEPs across both Kv1.3 blocker treatment groups, consistent with the different cellular and molecular effects of both Kv1.3 blockers in the brain, despite specific Kv1.3 blockade and shared reduction in Aβ pathology **(Fig. 7G)**. We also asked how CSF changes due to either Kv1.3 blocker, compared to changes of the same proteins in the brain proteome **(Supp. Fig. 6I & 6J)**. We found that following PAP-1 and ShK-223 treatment, there was little evidence for directionally concordant regulation between CSF and brain proteomes. To determine whether CSF proteins that reflect PAP-1 or ShK-223 treatment effects are relevant to human AD CSF biomarker changes, we cross-referenced PAP-1 and ShK-223 related CSF DEPs against CSF proteins that change in human AD^61^. This analysis showed that PAP-1 treatment reduced NEFL and RPH3A, both of which are increased with AD progression in human CSF. In contrast ShK-223 treatment reduced NME1 and GDA, which are increased with AD progression in human CSF **(Fig. 7H)**.

**Figure 7.**
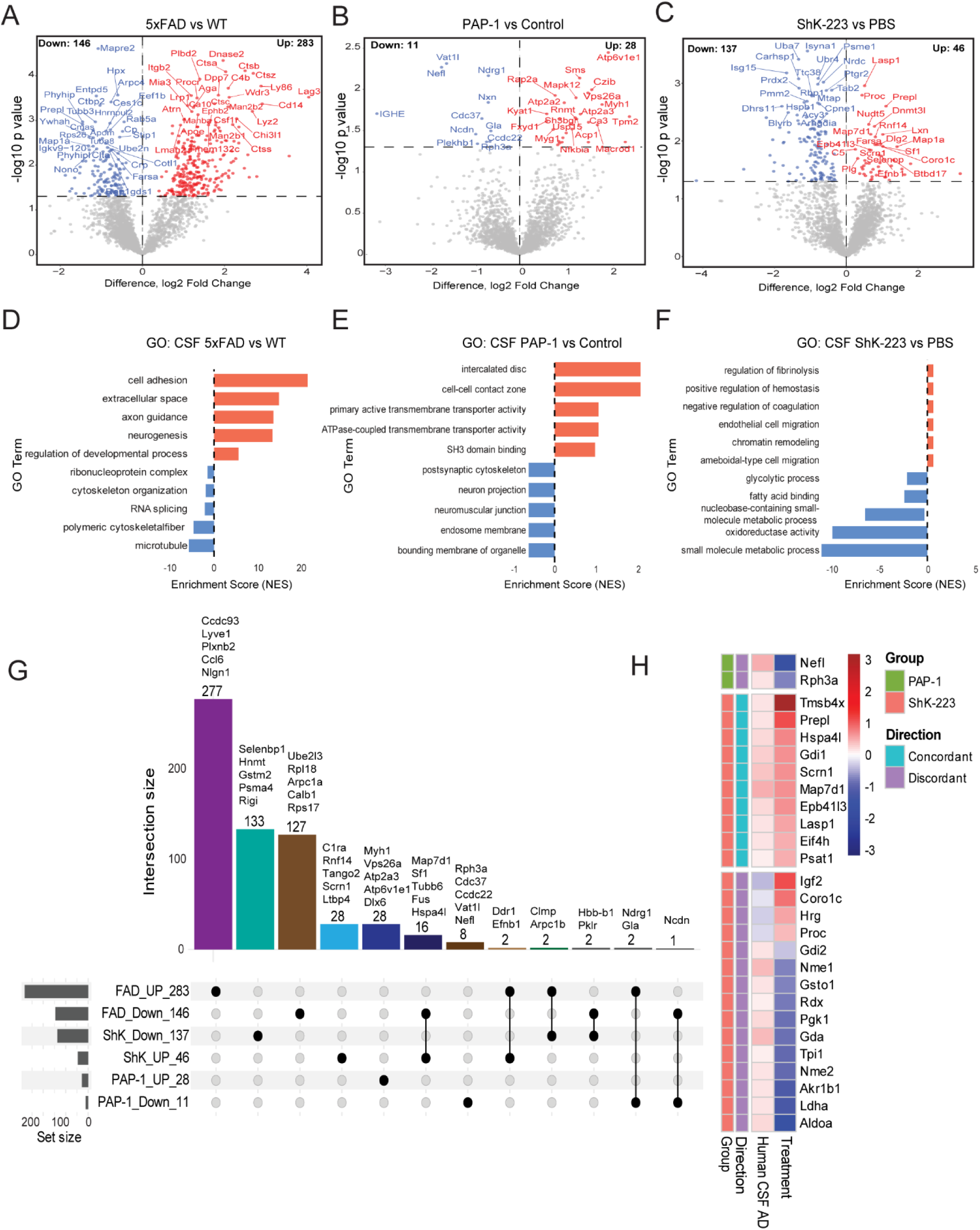
Proteomic alterations in mouse CSF following Kv1.3 inhibition with PAP-1 and ShK-223 and comparison with Alzheimer’s disease (AD) CSF signatures: **(A-C)** Volcano plots display differentially expressed CSF proteins in (**A**) 5xFAD vs WT (**B**) PAP-1 vs Control and (**C**) ShK-223vs PBS. Proteins with ≥ 0-fold change and p.value <0.05 are highlighted, with upregulated proteins in red and downregulated in blue. (**D-F**) Gene Ontology enrichment of significantly altered proteins from **(D**) 5xFAD vs WT, (**E**) PAP-1 vs Control, and (**F)** ShK-223 vs PBS. Bar plots show top biological processes associated with upregulated (red) and downregulated (blue) proteins. **(G)** UpSetR plot illustrates the intersection of significantly regulated CSF proteins across PAP-1, ShK-223, and 5xFAD significant upregulated and down regulated proteins intersection. Vertical bars indicate intersection size, while dot matrices denote shared or unique protein sets. Selected overlapping proteins are labeled, highlighting both conserved and condition-specific CSF signatures. (**H)** Concordant and discordant regulation of CSF proteins across mouse and human AD pathology: Heatmap displaying selected CSF proteins categorized as concordantly or discordantly regulated across PAP-1, ShK-223, and human AD CSF. Rows represent proteins and columns correspond to treatment, with the color scale indicating the direction and magnitude of log2 fold change. See Supp. Fig. 6 for related supplemental analyses and figures.

## DISCUSSION

AD pathophysiology involves complex interplay between different molecular pathways within and across cell types ultimately resulting in increased neuroinflammation, loss of neuronal function and synaptic as well as cellular degeneration^62^. In AD, higher levels of microglial Kv1.3 potassium channel have been associated with proinflammatory profiles of microglia^20^. Accordingly, inhibition of Kv1.3 by the small molecule PAP-1 or the peptide ShK-223 decreases AD-driven neuroinflammation and Aβ pathology in mouse models^21, 36^. Despite these common effects on Aβ pathology, it is unclear whether the highly brain-penetrant and cell membrane permeant PAP-1 and the less brain-penetrant and cell membrane impermeant ShK-223 have similar molecular and cellular effects in the brain. A large body of literature supports a role for Kv1.3 channels in microglia in AD pathology, although Kv1.3 transcript and protein have been detected in other cell types in the brain, as well as in immune cells outside the brain^30, 31, 32, 33^. This study presents snRNA seq and global transcriptomic as well as proteomic results for systematically uncovering different cellular responses and molecular pathways underlying the effectiveness of two Kv1.3 blockers in the 5xFAD mice. Concurrent with reduced Aβ pathology, these multi-omics data show that Kv1.3 blockade enhances molecular signatures of synaptic health, myelination, neurovascular coupling, and pro-resilience pathways, while decreasing reactive astrocytosis and neuroinflammation, showing broad neuroprotective mechanisms for both Kv1.3 blockers. One important anti-inflammatory effect of Kv1.3 blockade included decreased STAT1/3 phosphorylation in the brain. Using proximity-based proteomics and co-immunoprecipitation, we verified protein-protein interactions between Kv1.3 and STAT1 and STAT3 proteins in microglia. Our reproducible findings across cell systems using both pharmacological and genetic Kv1.3 manipulation in microglia, confirmed novel functional coupling between Kv1.3 channels and type 2 (but not type 1) IFN signaling. Based on these observed beneficial and neuro-immunomodulatory effects of Kv1.3 blockers and the emerging translational potential of Kv1.3 blockers in AD, we also identified novel biofluid protein biomarkers in CSF that reflect therapeutic effects of Kv1.3 blockade.

Our results show comparative efficacies of both Kv1.3 blockers (PAP-1 and ShK-223) in the 5xFAD mouse model using neuropathological, cellular and molecular approaches. Using standard biochemical and histological assays, we confirmed the decrease of Aβ plaques upon treatment with both PAP-1 and ShK-223, confirming findings from previous studies^8, 21, 36^. While blockade of Kv1.3 by PAP-1 reduces levels of total Aβ measured by ELISA of brain lysate, as well as Aβ plaque load in the cortex, ShK-223 reduced Aβ plaque load without effects on brain Aβ ELISA measures. This suggests that Kv1.3 may have differential effects on Aβ42 production and clearance mechanisms, which may also be related to different brain bioavailability and cellular targets of both molecules. Several snRNA seq studies have identified abnormal immune activation in microglia, impaired synaptic signaling in neurons, and myelination deficits in oligodendrocytes as the dysregulated pathways in AD mice models and AD patients ^63, 64, 65, 66^. To understand how these dysregulated pathways are affected by Kv1.3 blockade, we employed snRNA seq in a 5xFAD mice model treated with either PAP-1 or ShK-223. Our data provides a comprehensive understanding of cell type specific responses to Kv1.3 blockade and highlights several key molecules and pathways which contribute toward the overall neuroprotective effects of Kv1.3 blockade. Unexpectedly, we found oligodendrocytes to be the most transcriptionally responsive cell type with promyelination effects of both PAP-1 and ShK-223 treatment in 5xFAD mice. Myelin alterations have been shown to be associated with exacerbated amyloid deposition in both preclinical mouse models and AD patients^67, 68^. In a recent study, blockade of mitochondrial Kv1.3 also resulted in reduced demyelination by limiting activated microglia mediated neuroinflammation in mouse models of multiple sclerosis^69^. Given this, the promyelination effect of both Kv1.3 blockers on oligodendrocytes may augment myelin repair in AD-related demyelination, although this will require future investigation. Further, our snRNA seq findings from microglia upon PAP-1 treatment did not result in changes in proportions or DAM or homeostatic microglia, although several phagocytic and complement-related genes were altered by PAP-1 treatment. Maezawa *et al* have also shown that the Kv1.3 blockade using PAP-1in AD mice model inhibits microglial activation and enhances Aβ clearance capacity^36^. Together, these findings suggest that Kv1.3 blockers alter microglial profiles without a systematic change from DAM to homeostatic states. This possibility was also suggested by our prior work,^19, 21^ where ShK-223 skewed microglial gene signatures towards pro-phagocytic profiles that may be better adept at clearance of Aβ plaques. We also found that PAP-1 treatment in 5xFAD mice reduced astrogliosis, and both molecules altered the molecular profiles of astrocytes. Our bulk transcriptomic GO analysis also showed gliogenesis and glial cell differentiation as a downregulated term. These findings were further supported by reduced levels of GFAP protein abundance in our bulk proteomic analysis which was validated using western blot. Previous reports also suggest that functional deficits in AD mouse models can be improved by modulating the reactivity state of astrocytes^70^. While the effects of Kv1.3 blockade on microglia are predicted given existing roles for Kv1.3 in this cell type, our observed effects of both blockers on oligodendrocytes and astrocytes raise the possibility that Kv1.3 blockade has direct or indirect effects on these glial sub-populations. Future studies that will apply conditional cell type-specific genetic manipulation of Kv1.3 will help provide additional insights into cell type-specific contributions of Kv1.3 in AD pathology.

Consensus findings from our snRNA seq analyses as well as bulk transcriptomics and proteomics analyses of brain tissues identified conserved effects on cell types and cell-cell communications, some of which were shared by both Kv1.3 blockers. Among shared effects in glial cell types, our results nominated *Fth1* as being upregulated by both Kv1.3 blockers in microglia, astrocytes and oligodendrocytes. Intriguingly, the *Fth1* gene responsible for iron storage in a nontoxic form in the cytosol gets upregulated upon both PAP-1 and ShK-223 in all the glial cell types. Iron is a requirement for generation of myelin as demonstrated by dietary iron restricted animal studies^71^. However, free unbound iron capable of generating free radicals requires efficient antioxidant systems to protect neurons from oxidative damage due to their high oxygen requirements^72^. Recently, Mukherjee *et al* showed that *Fth1*, highly expressed by oligodendrocytes, is secreted via extracellular vesicles and plays an important role in protection against iron mediated ferroptotic axonal damage^73^. Though there are no reports of *Fth1* being secreted by microglia, *Fth1* levels are shown to be downregulated in microglia in AD^12^. PAP-1 mediated upregulation of *Fth1* can therefore represent a corrective measure. However, future validation and mechanistic studies are warranted to study the role of *Fth1* in glial cells in context of Kv1.3 blockade in AD models.

Our snRNA seq-based cell-cell communication analysis also showed an increased interaction between neurons, glial cells, and endothelial cells upon PAP-1 treatment in 5xFAD mice, suggesting an increase in neurovascular coupling. Interestingly, the term neurovascular coupling signaling pathway consistently identified in pathway analyses of both bulk brain transcriptomic and proteomic analyses. Moreover, our ligand receptor analysis in PAP-1 treated 5xFAD mice nominated several ligand receptor pairs like NRG1-ERBB4, PTN –PTPRZ1 and SLIT2-ROBO2 which are responsible for functional changes like restoration of E/I balance, and remyelination along with migration and polarization of endothelial^51, 53 74^. All these changes may contribute to the restorative effects of PAP-1. Additionally, the overall neuroprotective effects of Kv1.3 blockade by PAP-1 can also be explained by increased synaptogenesis and expression of pro-resilience proteins. Both synaptogenesis and synaptic membrane terms were consistently upregulated in IPA and GO analysis in our transcriptomic and proteomic data from PAP-1 treated 5xFAD mice. Moreover, we observed an increase in the protein abundance of PSD95 and SV2A in our PAP-1 treated 5xFAD mice bulk proteomic MS data further supporting the idea of increased synaptogenesis as a restorative mechanism for AD induced synaptic loss. Similarly, we observed a preferential effect of PAP-1 on increasing the pro resilience proteins including NRN1, the top pro-resilience protein identified in humans^75^. Strikingly, the upregulation of NRN1 is very consistent across our bulk proteomic, bulk transcriptomic, and pseudobulk analysis of glutamatergic neurons in PAP-1 treated group further signifying the importance of this protein in resilience pathways. Along with *Nrn1*, we also found the upregulation of another important determinant of neuronal vulnerability, Reelin, in our snRNA seq analysis from GABAergic neurons upon ShK-223 treatment suggesting ShK-223 also effects the resilience mechanisms for neuro protection^76^.

Our results also indicate effects of PAP-1, but not ShK-223, on mitochondrial proteins in AD pathology. In lieu of emerging roles for mitochondria-localized Kv1.3 channels in mammalian cells^33, 77^ brain mitochondrial effects of PAP-1 may therefore involve direct modulation of mitochondrial Kv1.3 channels by the ability of PAP-1 to enter the cell, while ShK-223 is cell-impermeant. Only PAP-1 treatment and not ShK-223 increased the mitochondrial protein levels compared to control, including the mitochondrial outer membrane transport protein TOMM20, which we also validated by western blot. The global effects of PAP-1 on mitochondrial proteins could be explained by regulation of mitochondrial biogenesis, reduced mitophagy, or disrupted mitochondrial transport. Interestingly, the mitochondrial division signaling pathway was also upregulated in our pathway analyses of PAP-1 treated bulk brain proteomics data, which can be a result of increased cellular energy demand or stress. Recently, the Kv1.3 blockade using PAP-1 in a robust mouse model of ALS showed an improved mitochondrial morphology and function suggesting that Kv1.3 activity is implicated in degeneration of mitochondria^78^. The compartment specific effect of PAP-1 was also shown in our *in vitro* experiment using BV2 microglia cells. We found that only PAP-1, but not ShK-223, inhibited IFN-γ induced phosphorylation of STAT proteins. Previously, mitochondrial complex 1 has been shown to be a positive regulator of IFN-γ response^79^. Moreover, we also show physical interaction between the STAT1/3 proteins and Kv1.3, specifically STAT3 and Kv1.3 interaction has been reported by others using BioID approach^80^. Since Kv1.3 interacts with and regulates complex 1 of the electron transport chain in melanoma cells, we speculate this can be the mechanism of preferential inhibition of IFN-γ induced phosphorylation of STAT proteins^81^. Taken together, these findings provide novel insights into regulation of ferroptosis, neurovascular coupling and mitochondria by Kv1.3 blockers in AD pathology.

From a translational perspective we also identified novel protein biomarkers in CSF that reflect efficacy of Kv1.3 blockade in AD pathology. Neurofilament-light (NEFL) and excitatory neuronal and cognitive resilience protein RPH3A, both were both decreased in the CSF upon PAP-1 treatment. Both these proteins are also increased in human AD CSF as compared to controls. NEFL reflects axonal integrity and levels in CSF and plasma positively correlate with level of neurodegeneration or axonal injury, across several neurological diseases^82^. RPH3A is another promising AD-relevant CSF biomarker that changes with Kv1.3 blockade. RPH3A is a synaptic protein that is highly expressed by excitatory neurons which decrease in the brain but increase in AD CSF^83, 84^. As compared to CSF changes due to PAP-1 treatment, we observed stronger effects of ShK-223 treatment, including reduced levels of ISG15, a major immune protein that is upregulated due to interferon signaling^85^. Among CSF biomarkers of ShK-223 treatment effect, 11 were increased in human AD and decreased by ShK-223, while 4 were decreased in human AD but increased by ShK-223, representing a group of proteins reflecting disease-modification due to Kv1.3 blockade. We suspect that the minimal overlap between CSF biomarkers of treatment effect of PAP-1 and ShK-223 are due to different cellular targets and differences in brain bioavailability of both drugs, as also evidenced by our brain-level proteomics and transcriptomics results. Beyond CSF, future studies should further assess these proteins in larger cohorts as well as in the plasma. In the era of Abeta-directed therapies for AD, biofluid protein biomarkers of drug efficacy can be very useful in early phase human trials, to ensure target engagement.

One of the major limitations of the current study is that we have not assessed sex specific effects of Kv1.3 blockade. Female 5xFAD are known to show increased Aβ pathology and molecular signatures of AD compared to male mice and hence we tested the efficacy of Kv1.3 blockers in these mice ^86^. To gain a comprehensive sex specific effect of Kv1.3 blockade, studies need to be performed in male mice separately. Secondly, additional biological replicates are needed for conclusive findings in behavior studies using PAP-1 and ShK-223. Lastly, in our pseudobulk analysis, the number of nuclei sequenced for low abundance cells was small, which can affect the overall accuracy of the analysis. Also, in our pseudobulk analysis, we had some unexpectedly profound effects of Kv1.3 blockade on oligodendrocytes, neurons, and astrocytes other than microglia where channel presence is extensively studied. In future studies, the mechanistic basis of these cell type specific effects needs to be further validated. It will also be interesting to test the efficacy of these pharmacological inhibitors in other mouse models of AD like Tau model. In summary, our results demonstrate that long term blockade of Kv1.3 in an amyloid mice model of AD has beneficial neuroprotective outcome along with inhibition of neuroinflammatory pathways and augmentation of pro-resilience pathways. These findings further strengthen the rationale to pursue the Kv1.3 channel as a putative drug target in AD with more potent and selective brain permeant inhibitors targeting the ion channel.

## Supporting information

Table 1 related to fig 1

Table 4 related to fig 4

Table 5 related to fig 5

Table 6 related to fig 6

Table 7 related to fig 7

Table 2 related to fig 2

Table 3 related to fig 3

Supp. table 1 related to supp fig 1

Supp. table2 related to supp fig 2

Supp. table3 related to supp fig 3

Supp. table4 related to supp fig 4

Supp. table5 related to supp fig 5

Supp. table6 related to supp fig 6

Supplemental datasheet 7 for materials and antibodies

table legends for tables and supp. tables

## Abbreviations

Aβ: Amyloid beta
AD: Alzheimer’s Disease
AMPA: α-amino-3-hydroxy-5-methyl-4-isoxazolepropionic acid
BP: Biological process
CC: Cellular component
CNS: Central nervous system
CREB: cAMP-response element binding protein
CSF: Cerebrospinal fluid
DAMs: Disease-associated microglia
DEG: Differentially expressed genes
DIA: Data-independent acquisition
ELISA: Enzyme-linked immunosorbent assay
GABA: Gamma-aminobutyric acid
GFAP: Glial fibrillary acidic protein GO Gene ontology
IBA1: Allograft inflammatory factor 1
IF: Immunofluorescence
IFN: Interferon
IFN-α: Interferon alpha
IFN-β: Interferon beta
IFN-γ: Interferon gamma
IPA: Ingenuity pathway analysis
i.p: Intraperitoneal
IRMs: Interferon-responsive microglia
JAK: Janus kinase
JAK1: Janus kinase 1
KO: Knockout
Kv1.3: Potassium voltage-gated channel, shaker-related subfamily, member 3
L/R: Ligand–receptor
LR: Ligand and receptors
MA: Bland–Altman plot
MF: Molecular function min-Minute
mRNA: Messenger RNA
MS: Mass spectrometry
NGF: Nerve growth factor
NMDAR: N-methyl-D-aspartate receptor
NRN1: Neuritin 1
OPCs: Oligodendrocyte progenitor cells
PAP-1: 5-(4-Phenoxybutoxy) psoralen
PD: Parkinson’s Disease
PRRs: Pathogen recognition receptors
PSD-95: Postsynaptic density protein 95
QC: Quality control
RNA: Ribonucleic acid
RNA seq: RNA sequencing
ROS: Reactive oxygen species
S: second
ShK-223: Stichodactyla toxin 223
snRNA-seq: Single-nucleus RNA sequencing
STAT: Signal transducer and activator of transcription
STAT1: Signal transducer and activator of transcription 1
STAT3: Signal transducer and activator of transcription 3
SV2A: Synaptic vesicle glycoprotein 2A
TEM: T-effector memory cells
TOM: Translocase of the outer membrane
TOMM20: Mitochondrial import receptor subunit TOM20 homolog
UMAP: Uniform Manifold Approximation and Projection
UMIs: Unique molecular identifiers
WNT: Wingless Int-1
WT: Wild type

## METHODS

### Reagents

A detailed list of reagents used in these studies, including antibodies, is provided in **the Supplemental Datasheet 7**

### Cell culture

BV2 microglia transduced with V5-TurboID-Kv1.3 fusion construct termed as BV-2 Sr73^27^ or MMC cells were cultured in DMEM-F12 (10% FBS and 1% Pen/Strep) and plated in a 6-well plate at 1 million cells and were allowed to adhere to the plate 24h prior to experiments. Cells were treated with PAP-1 (1 µM) or ShK-223 (20nM) for 30 mins for Kv1.3 channel blockade followed by exposure to either IFN α (100 U/ml) or IFN γ (10 ng/ml) for 30 mins. Supernatant was then removed, and cells were washed with chilled PBS complemented with protease and phosphatase inhibitor. Cells were lysed in the specific lysis buffer depending on the assays to be performed.

Primary microglia were isolated from 1–3-day-old pups born to C57BL/6J wild-type or *Kcna3* knockout mice. Briefly, the *Kcna3* gene was modified by inclusion of loxp sites 5’ upstream and 3’ downstream of the *Kcna3* exon, including a neomycin selection cassette. Mice were then serially back-crossed for 8 generations onto the C57BL/6J background. Subsequently, we crossed *Kcna3*-floxed mice with CMV-Cre mice, we derived homozygous *Kcna3*-KO mice, which were then confirmed by *Kcna3* genotyping. These mice were bred as homozygotes, along with WT counterparts, and were used to generate p0-p1 pups from which primary microglia cultures were derived. Further details about this *Kcna3* KO mouse line and conditional *Kcna3*-floxed line, will be published separately. C57BL/6J wild-type or *Kcna3* knockout mice pups were deeply anesthetized and euthanized by cooling on ice for 3-4 minutes, followed immediately by decapitation. Brains were carefully removed from the skull without contaminating the brain tissue and placed in 2 mL ice-cold 1× PBS in a 30 mm cell culture dish. Using sterilized fine surgical forceps and a dissection microscope, the cortices were isolated by carefully removing the meningeal tissue. Cortical tissue was then transferred into a 15 mL centrifuge tube containing 1 mL ice-cold 1× DMEM/F-12 and transported to the cell culture hood. The tissue was washed with pre-warmed DMEM/F-12 and transferred into a 12-well plate. Cortices from 3–4 mice were placed in each well and chopped into small pieces using a scalpel blade without mixing genotypes. Enzymatic digestion was performed by adding by adding 500 µl of 0.5% Collagenase Type II and 100 mg/mL Dispase II – Neutral Protease in ratio 1:1, along with 75 µL of 1 mg/mL DNase I, under constant agitation (150 rpm) for 45 min at 37°C in a 5% CO₂ incubator. Following digestion, the tissue was mechanically dissociated using fire-polished Pasteur pipettes (3–5 passes) until the suspension became homogeneous. Cells were filtered through a 40 µm cell strainer into a 50 mL conical tube and centrifuged at 1,000 × *g* for 5 min at 4°C. The supernatant was carefully discarded, and the cell pellet was resuspended in pre-warmed DMEM/F-12 medium. Cells were plated in poly-L-lysine–coated 150 cm² culture flasks and incubated at 37°C in a 5% CO₂ incubator. After two days, cells were washed with 1× PBS and fresh medium was added. Cultures were maintained until confluence (between 2-3 weeks), with medium changes every 2–3 days.

The passaging and purification of microglia was performed according to a previously published protocol with a slight modification.^87^. In brief, cells were first washed with cold PBS, and 8 mL of 0.05% trypsin (diluted in DMEM/F-12) was added. Cells were incubated for 35–40 min, until the astrocyte layer began to detach. The astrocytic layer was discarded, and 4 mL of serum-containing medium was added to wash the flask. Cells were again washed with ice-cold PBS to remove any residual floating cells. To collect adherent microglial cells, 7 mL of TrypLE Express was added and incubated for 8 min. The reaction was stopped by adding 5 mL of DMEM/F-12 medium. Cells which were still attached to the surface were scraped and all the cells were collected into a 50 mL centrifuge tube, centrifuged at 1000 × *g* for 5 min, and resuspended in 5 mL fresh DMEM/F-12 medium. Microglia were seeded at a density of 1 × 10⁶ cells per well in poly-L-lysine (100 µg/ml) pre-coated 6-well plates.

### Animals and drug administration

6 and 9-month-old female mice were used for all experiments. Animals were housed in the Department of Animal Resources at Emory University under a 12 h light/12 h dark cycle with ad libitum access to food and water and kept under environmentally controlled and pathogen-free conditions. All experiments involving animal procedures were approved by the Emory University Institutional Animal Care and Use Committee. The mice were aged 3 months and 6 months. Mice were then treated with ShK-223, PBS, or PAP-1. ShK-223 and PBS were injected twice a week at 100 μg/kg/dose (2 mg/mL in PBS with 0.1% BSA). PAP-1 was administered through food chow at 1100 ppm. Chow was prepared by PMI Nutrition International using the 5001 Lab Diet.

### Behavioral studies

Following three months of treatment, behavioral analysis was performed. Treatment was continued during behavioral analysis for the final week of treatment.

#### Water Maze

Mouse Morris Water Maze training takes place in a round, water-filled tub (52-inch diameter) in an environment rich with extra maze cues and a small platform 1 cm below the surface. White tempera paint (non-toxic) was added to the water to make it opaque so that the mice could see the platform. Mice were placed in the water maze with their paws touching the wall from 4 different starting positions (N, S, E, W) in water that starts at 25 °C and typically declines to 22 °C by the time a whole group of mice is tested. An invisible escape platform was in the same spatial location 1 cm below the water surface, independent of a subject’s start position on a particular trial. In this manner, subjects utilized extra maze cues to determine the platform’s location. Each subject was given 4 trials/day for 5 days with a 15-min inter-trial interval. The maximum trial length was 60 seconds(s) and if subjects did not reach the platform in the allotted time, they were manually guided to it. Upon reaching the invisible escape platform, subjects were left on it for an additional 5 s to allow for survey of the spatial cues in the environment to guide future navigation to the platform. After each trial, subjects were kept in a dry plastic holding cage filled with paper towels to allow them time to dry off. Following the 5 days of task acquisition, a probe trial was presented during which time the platform was removed, and the amount of time and distance swam in the quadrant which previously contained the escape platform during task acquisition was measured over 60 s. All trials were videotaped and performance analyzed by MazeScan (Clever Sys, Inc.), the output of which was latency, distance, and speed. For each day, the average of the four trial sessions was provided of latency, distance, and speed. This was then graphed and analyzed using Prism 10 (GraphPad). Anova was utilized to calculate changes between groups on a given day.

#### Fear Conditioning

Cued and contextual conditioning is a fear conditioning task which measures the ability of a rodent to form and retain an association between an aversive experience and environmental cues. The Emory Behavior core used a standard fear conditioning paradigm over three days. Mice were placed in the fear conditioning apparatus (7” W, 7” D X 12” H, Coulbourn) composed of Plexiglas with a metal shock grid floor and allowed to explore the enclosure for 3 min. Following this habituation period, 3 conditioned stimulus (CS)-unconditioned stimulus (US) pairings were presented with a 1 min intertrial interval. The CS consisted of a 20 S 85 Decibel (dB) tone, and the US consisted of 2 seconds of a 0.5 mA foot shock, which co-terminated with each CS presentation. This level of foot shock is modest; by comparison, stress-induced reinstatement of drug seeking uses 15 min of intermittent 0.8 mA foot shock. Shock was delivered via a Precision Animal Shocker (Colbourn) connected to each fear-conditioning chamber. The shock level was set and verified prior to each training session. The voltage varied between 4-5 volts because the system uses resistance-regulated constant-current output. One minute following the last CS-US presentation, animals were returned to their home cage. On day 2, the animals were presented with a context test, during which subjects were placed in the same chamber used during conditioning on Day 1, and the amount of freezing was recorded via a camera and the software provided by Colbourn. No shocks were given during the context test. On day 3, a tone test was presented, during which time subjects were exposed to the CS in a novel compartment. Initially, animals could explore the novel context for 2 min. Following this habituation period, the 85 dB was presented for 6-min, and the amount of freezing behavior was recorded. No shocks were given during the tone test. The mice were returned after three days when the testing was complete, but we maintained chow and IP treatment during that time. The percentage freeze was compared to seconds in prism for the training day, the context, and the cue in Prism 10 (GraphPad). One way Anova was utilized to evaluate the significance of groups at different time points. (n=3 or 6)

### CSF collections from mouse cisterna magna

The mice were anaesthetized by intraperitoneal injection of a mixture of ketamine (73.5 mg/kg; Akron, USA), xylazine (9.2 mg/kg; Bayer Pharma, Germany), and acepromazine maleate (2.75 mg/kg; Boehringer Ingelheim, USA) in 200 µL of 0.9% (v/w) NaCl. After the withdrawal of sensory reflexes, mice were shaved on the back of the neck to the base of the skull and then disinfected with ethanol wipes. The mouse’s head was stabilized on a stereotaxic frame (David Kopf Instruments, Tujunga, CA, USA) without fixing the jaw on the tooth adapter. No lateral head movement ensured proper head fixation on the stereotactic frame. To level the mouse on the stereotaxic frame, the body was placed on a custom-made box. A linear skin incision was made in the midline from the top of the skull to the neck using a scalpel blade (No.11). The skin was retracted, and the muscles were exposed after the removal of the superficial layer of connective tissue using forceps. Using fine scissors muscles were detached from the base of the occipital bone. Extra muscles and blood were removed using a cotton tip, and the area was cleaned for better visualization of the Cisterna Magna. Little connective tissue was left on the top of the dura that stabilized the Hamilton needle and prevented the backflow of the CSF. Hemostasis was prevented during the entire process, either using a cotton swab or a suction aspirator. A Hamilton syringe containing a 30 G needle was attached to a micro syringe injector pump (KD Scientific) and placed on a stereotaxic frame. The Hamilton syringe needle was adjusted to a 60-degree angle. Mice heads were pressed down using a tooth bar adaptor to expose the cisterna magna syringe injector needle was aligned with the cisterna magna linearly, and the needle was inserted into the cisterna magna after puncturing the dura. CSF was drawn into the syringe by slow and smooth aspiration by starting the microinjector pump aspirator at a flow rate of 8 µl/min. CSF was inspected with the naked eye for quality and clarity. The CSF was transferred into a new 500 µl centrifuge tube and briefly spun down, and the supernatant was transferred into a new vial and snap frozen on dry ice for further analysis.

### Tissue collection

After CSF collections, mice were transcardially perfused with ice-cold PBS (1X). Mice brains were immediately removed by dissection without damaging the brain structure. The right hemisphere of the brain was snap frozen on dry ice either intact or after regional dissection (cerebellum, brain stem, cortex, hippocampus, and striatum/thalamus) for downstream proteomics analysis, while the left hemisphere was post-fixed in 4% PFA for 24 hours and then transferred to the 30% sucrose for immunohistological studies.

### Tissue processing for protein-based analysis, including Western blot

Frozen brain tissues without thawing, was weighed and added to 1.5 mL Rino tubes (TUBE1R5-S; Next Advance) containing a small scoop of stainless-steel beads (0.9–2 mm in diameter, SSB14B; Next Advance), urea lysis buffer (8 M urea, 10 mM Tris, 100 mM NaH2PO4, pH 8.5; five volumes of the tissue weight) and 1X HALT protease inhibitor cocktail without EDTA. Tissues were homogenized in a Bullet Blender (Next Advance) twice for 5 min cycles at 4 °C at a speed setting of 10. Tissues were further sonicated, consisting of 5 s of active sonication at 30% amplitude with 5 s incubation periods on ice cold water. Homogenates were allowed to sit for 5 min on ice and then centrifuged for 5 min at 12000-G, and the supernatants were transferred to a new tube. Protein concentration was determined by BCA assay using Pierce™ BCA Protein Assay Kit. To evaluate protein of interest 15 μg of protein was added to 4× Laemmli buffer supplemented with β-mercaptoethanol (1:10). This was then boiled for 10 min at 95^°^C and centrifuged for 10 min at 13000 RPM. Samples were resolved on a BOLT 4%–12% Bis–Tris gel along with a molecular weight indicating ladder. Protein was transferred to either a nitrocellulose or polyvinylidene difluoride membrane using the iBLOT2 stack system. The membrane was blocked for 1h at room temperature using the StartingBlock blocking buffer. Blots were incubated with various primary antibodies overnight at 4^°^C with slight agitation. Blots were then washed with TBS-T (0.1%) 3 times, 10 minutes each. Secondary antibody incubation was performed for 1h at room temperature with agitation. The membrane was imaged using the Odyssey Li-COR system or Bio-Rad ChemiDoc MP Imaging system. Densitometry analyses for band density were done using ImageJ software and statistical significance was measured using T-test on Prism 10 software.

### Immunohistochemistry and image quantification

Brain tissue was fixed in 4% PFA for 24h then transitioned to 30% sucrose. Fixed brain tissue was then sliced, sagittally, using a cryostat (Lyca Biosystem # 1950) at 40 µm thickness and transferred into the cryoprotectant liquid and stored at -20*C until further use. For staining, free-floating sections were washed 3 times in TBS to get rid of any cryoprotectant. The sections then were permeabilized and blocked using 5% horse serum in TBS-T (0.3 % Triton X100 in TBS) for 1 h. The sections were further incubated with desired primary antibodies in TBS-T buffer containing 1% serum, overnight at 4* C on a shaker. After primary antibody incubation, tissues were washed 3x with 1x TBS for 5 min each and incubated with respective secondary antibodies at dilution (1:500). for 1h. Nuclear staining was then performed with DAPI (<u>1 µg/ml</u>) during the first washing step. After three washes with1xTBS for 5 min, stained tissue sections were mounted on a glass slide and left to air dry. Tissues were covered using Prolong Diamond mounting media and kept in the fridge until they were imaged.

Images were captured using a Keyence BZ-X810 microscope with 4x, 20x, and 40x objective lenses. Image processing was performed either using the Keyence BZ-X810 Analyzer or Image J software (FIJI Version 1.53). Aβ plaque size and % area was measured using the Keyence image analysis software. The average for three slices per mouse (n=3) was calculated for the subiculum and cortex and graphed utilizing Prism (GraphPad, 10.4.2). Statistical significance was evaluated utilizing an unpaired t-test.

### Amyloid-β42 ELISA

ELISA was performed by using Human Aβ (aa1-42) Quantikine ELISA Kit. In brief, after the accurate measurement of proteins, the concentration was adjusted across the samples, and ELISA was performed on 0.3 µg of the protein for each sample in duplicate according to the manufacturer’s protocol The colorimetric assay was run on a Tecan Infinite pro200 at 450 nm, and Amyloid-β abundance was matched to a standard curve. Total Amyloid-β abundance was graphed using Prism (Graphpad, 10.4.2). Statistical significance was evaluated utilizing an unpaired t-test (n=3 or 6).

### Preparation of Samples for snRNA seq

Flash frozen brain tissue (20 mg) from mice was utilized for nuclear isolation adapted from Corces et al. ^88, 89^ In brief, tissue was homogenized in a homogenization buffer (HB) consisting of 1 M sucrose, 1 M KCl, 1 M Mg2Cl, 1 M Tricine-KOH (pH 7.8), 1 M DTT, 500 mM spermidine, 150 mM spermine, 10% NP-40, 0.5 tablets of cOmplete protease inhibitor, a 1:100 dilution of Ribolock, 5 mg/mL Antimycin D, 10 mM triptolide, and 10 mg/mL Anisomycin on ice. Completion of lysis was confirmed using a hemocytometer by visualizationof breakdown of cell membrane while maintaining nuclear membrane. Lysates were filtered with a 40 μm strainer then centrifuged at 600 x g for 7 min at 4 °C, Cell pellets were resuspended in HB and mixed with a 50% Iodixanol solution (diluted in HB) at a 1:1 ratio. A layer of 30% Iodixanol solution (diluted in HB) was layered under the 25% mixture and a 40% Iodixanol solution (diluted in HB) was layered under the 30% solution. This was then centrifuged for 20 min at 4 °C at 3,000 RCF with the brake turned off. Nuclei were located at the 30% -40 % interface and collected using a pipet. Nuclei were then resuspended in wash buffer (WB) consisting of 30 % bovine serum albumin (BSA), a 1:100 dilution of Ribolock, 5 mg/mL Antimycin D, 10 mM triptolide, and 10 mg/mL Anisomycin and centrifuged at 600xg for 10 min at 4 °C. Nuclei were counted on a hemocytometer then centrifuged at 600xg for 10 min at 4 °C and stored in 3mL of WB. Following immediately, 10,000 nuclei were prepared for Cell Multiplexing Oligo Labeling. Libraries were prepared using the Chromium Next GEM Single Cell 3’Reagent Kits v3.1. Libraries were evaluated via Bioanalyzer Libraries were then sequenced at 50,000 read pairs per nucleus using 2× 150 bp reads on an Illumina Novaseq 6000 instrument through Admera Health. Sequencing. data was processed using the 10x Genomics Cell Ranger ARC (cellranger-arc-2.0.0) pipeline with default parameters and aligned to the hg38 reference genome (refdata-cellranger-arc-GRCh38-2020-A-2.0.0).

### snRNA seq data processing and quality control

The CellRanger (v3.0.2 10X Genomics) software was employed to align the sequences and quantify gene expressions. We used the mouse reference genome mm10 to prepare a pre-mRNA reference according to the steps detailed by 10X Genomics. snRNA-seq libraries capture reads from both unspliced pre-messenger RNAs (mRNAs) and mature mRNAs; we first generated a pre-mRNA reference genome according to the instructions provided by 10× Genomics. We subsequently aligned the demultiplexed FASTQ files from Novogene to the refdata-gex-mm10-2020-A (mm10) pre-mRNA reference genome using CellRanger (version 3.0.1) with the default settings (G. X. Y. Zheng et al., 2017). Secondary filtering in Seurat (v3.0) removed low-quality nuclei with ≤300 detected genes, ≥10,000 UMIs, or ≥10% mitochondrial transcripts, yielding a final dataset of 137,375 nuclei and 28,608 genes. To address sample variability and condition-driven batch effects, we used the Harmony R package to remove the batch effect. For integrative analysis, log-normalized data were processed in Seurat by identifying 2,000 variable features per sample, followed by anchor-based integration (dims 1:30), scaling, and PCA. Clustering was performed with the first 30 PCs at clustering resolution 0.5, and visualized using UMAP, resulting in 24 clusters. Differentially expressed genes were identified with the Wilcoxon rank-sum test (log2FC ≥0.25, adjusted P <0.05), and clusters were annotated based on canonical marker expression and comparison to reference datasets (Tasic et al., 2016). Annotation accuracy was further confirmed using prediction scores from scCustomize R package (Marsh SE et al., 2021).

### Pseudobulk differential expression and pathway enrichment

For each cell type and experimental comparison, differential gene expression was analyzed using the glmGamPoi R package where raw count matrices were obtained from the *SCT* assay of seurat objects, and genes were retained if expressed in at least 10% of cells in both groups. The filtered matrices were normalized using deconvolution-based size factors, and generalized linear models with a gamma poisson distribution were fitted with glm_gp. Differential expressions were assessed with test_de function by specifying the relevant contrasts as PAP-1 versus control, ShK-223 versus PBS. Genes with an absolute log fold-change of ≥0.25 and adjusted *p* < 0.05 were considered significantly regulated and classified as upregulated or downregulated. The analysis was conducted separately for each cell type to capture specific transcriptional changes. Gene Ontology enrichment was conducted using enrichR and ClusterProfiler. The top enriched pathways were visualized in R using ggplot2, highlighting functional differences across conditions and cell types.

### Cell-cell communication analysis

Cell to cell communication networks were inferred using CellChat v2 (Jin et al., 2021) with the built-in mouse ligand receptor database (*CellChatDB.mouse*). For this analysis, we used normalized count expressions from snRNAseq seurat object and cluster annotations were used as input, and standard preprocessing steps were applied. Differential communication between PAP-1 versus control and ShK-223 versus PBS was assessed by computing interaction probabilities with compute CommunProb and testing significance by permutation. Pathway level differences were quantified with compare Interactions and visualized using net visual bubble. Network centrality metrics were used to define major sending and receiving populations, and enriched ligand receptor interactions.

### RNA sample preparation and quality control of 5xFAD mouse brain

Total RNA was obtained from Admera Health. All samples were quality-checked using Agilent TapeStation, and only those with RNA Integrity Numbers (RIN) above 8.0 were included for sequencing. RNA concentrations ranged from ∼295 to 597 ng/µl with total yields between ∼11-19 µg.

### Differential expression and visualization

Paired-end RNA-seq reads were aligned to the Mus-musculus reference genome (GRCm39/mm10) using STAR (v2.7.11a) with gene annotations from the corresponding FASTA files provided by Admera Health. Transcript level protein coding counts generated by STAR were imported into R (v4.3.2) and analyzed with DESeq2 (v1.40.2). Differentially expressed genes were defined as those with adjusted *p* value < 0.05 (Benjamini Hochberg correction) and |log2 fold change| ≥ 0.25. Visualization of differential expression results was performed using the DESeq2 MA plot function and further customized with ggplot2 (v3.4.4).

### Protein digestion MS, protein identification, and quantification

#### S-trap digestion

Each sample was digested in an S-trap micro spin column (Protifi, USA) according to the manufacturer’s instructions. For each sample 50 µg of protein lysates were used. The samples were reconstituted in 5% SDS in 50 mM TEAB. The proteins were then reduced with 10 mM of dithiothreitol (DTT) at room temperature for 1 hour. The proteins were alkylated with 20 mM of iodoacetamide (IAA) in the dark for 30 mins. The alkylated proteins were acidified by adding phosphoric acid to a final concentration of 1.2% followed by addition of six volumes of binding/washing buffer (90% methanol; 100 mM TEAB; pH 7.5). The protein solution was loaded onto the S-trap filter in 70µl increments, repeated till all the samples were loaded onto the column at 4000g for 30s. The samples were then washed 3 times with binding/washing buffer in 150 µl increments. Membrane bound proteins were digested with 0.5 µg of lysyl (Lys-C) endopeptidase in 25 µl at 37°C for overnight. Then 1 ug of trypsin was added in 25 µl and incubated overnight at 37°C. The peptides were eluted stepwise with three elution buffers at volume of 80 µl, including 50 mM TEAB in water, 0.2% formic acid in water, and 50% acetonitrile in water. The pooled peptide solution was dried using speed-vacuum and dissolved in 20 µl of 0.1% formic acid for fluorescence-based peptide BCA.

#### DIA MS

Samples for DIA were analyzed on an LC–MS/MS consisting of a Vanquish Neo UHPLC (Thermo Fisher Scientific) nanoflow liquid chromatography system and an Orbitrap Exploris 480 mass spectrometer (Thermo Fisher Scientific) equipped with an EASY-Spray ion source. The peptides were reconstituted in 0.1% formic acid and loaded onto a trap column (300 μm inner diameters, 5 mm length, 5 μm C18 particles, PepMap Neo Trap, Thermo Fisher Scientific) before being separated on an EASY-spray HPLC analytical column (75 μm inner diameter, 60 cm length, 1.7 μm C18 particles, Aurora Frontier, Ionopticks) at a flow rate of 300 nL/minute. Mobile phase A and B were composed of HPLC water with 0.1% formic acid (FA), and 80% HPLC acetonitrile with 0.1% FA, respectively. The peptides were resolved by changing the gradient of mobile phase B as follows: 1% to 4% in 10 min, 4% to 25% in 60 min, and 25% to 65% in 45 min. The gradient was followed by a 20-minute wash step (65% to 99% in 1 min, hold at 99% for 19 min) and column re-equilibration. An MS analysis was performed in data-independent acquisition (DIA) mode using an m/z Range window mode. The voltage for electrospray ionization (ESI) was set to 2,100 V, and the ion transfer tube temperature was set to 275 °C. The DIA method consisted of one full MS1 scan followed by 60 DIA (MS2) scans. The MS1 scan range was set to 400–1000 m/z. MS1 scans were acquired at a resolution of 120,000 (at 200 m/z) with an RF Lens of 50%, a normalized AGC target of 300% (absolute target 3,000,000), and an ’Auto’ maximum injection time. The 60 subsequent DIA (MS2) scans were acquired at a resolution of 30,000, with a normalized AGC target of 1000% (absolute target 1,000,000) and an ’Auto’ maximum injection time. Fragmentation was induced by high-energy collision dissociation (HCD) at 28% of the normalized collision energy. The isolation window for precursor ions was set to 10 m/z with 1 m/z overlap, covering the precursor range m/z 400–1000. Internal calibration was carried out using the lock masses of plydimethylcyclosiloxane ions (m/z 445.12003) produced from ambient air.

#### Proteomic Data Processing and Quantification

The acquired DIA raw files were processed using DIA-NN (v2.2.0, Academia) in library-free mode. The search was conducted against a Mus musculus (Mouse) FASTA database (UniProt UP000000589, downloaded 2025.04.11) supplemented with common contaminants. Search parameters were set to Trypsin/P specificity, allowing for up to two missed cleavages. Carbamidomethylation (C) was set as a fixed modification. Variable modifications included N-terminal M excision, Oxidation (M), and N-terminal acetylation, with a maximum of three variable modifications per peptide. The peptide length was restricted to 7–30 amino acids, and the precursor charge range was set from 1 to 4. The analysis matched precursors within the m/z 400–1000 range and fragments within the m/z 150–2000 range. Mass tolerances for both MS1 and MS2 were set to 10 ppm, with a scan window of 15. The analysis used neural network (NN)-based, cross-validated machine learning and proteoform-level scoring. All results were filtered at a 1% precursor and protein group FDR. Protein quantification was performed using the QuantUMS strategy. Cross-run normalization was disabled within DIA-NN.

### LC-MS MS data analysis of 5xFAD mouse brain

Proteomic data from 5xFAD mouse brains were processed in Perseus (http://www.perseus-framework.org/). Proteins were retained if at least 50% of values were present in one group, and intensities were log2-transformed. Missing values were imputed in Perseus by random sampling from a normal distribution. To correct sample loading differences, column-sum median normalization was performed in R, where total protein intensity was calculated per sample, and each was scaled by the ratio of the global median column sum to its own sum. Group comparisons were assessed using one-way ANOVA and two-sample *t*-tests. Multiple testing correction was applied with the Benjamini Hochberg procedure (FDR < 0.05), with statistical significance defined at *p* < 0.05. Differentially expressed proteins were visualized using volcano plots generated in ggplot2, displaying log2 fold change against P-value.

### LC-MS MS data analysis of 5xFAD mouse CSF

Using a DIA mass spectrometry approach, CSF proteomics was executed out, finding approximately 6,000 protein groups. 2,495 proteins remained after proteins with more than 75% missing values were removed. Using a left-shifted gaussian distribution and log₂ transformation, missing data were imputed in Perseus software. A carefully selected panel of blood-associated proteins was used to evaluate blood contamination. As per sample, blood score was then calculated and regressed to account for blood-related volatility. Data were adjusted using column-sum median normalization method after blood correction. Differential protein expression was assessed using one-way ANOVA statistical testing, and nominal p-value < 0.05 was considered significant. In each comparison (PAP-1 against control, ShK-223 vs PBS, and 5xFAD vs WT), proteins with log2 fold change > 0 were considered upregulated, whereas those with log2 fold change < 0 were considered downregulated.

### GO enrichment analysis

Gene Ontology enrichment was performed using the cluster Profiler (enrichGO) R package with org.Mm.eg.db for mouse gene annotation, querying all three ontologies: Biological Process (BP), Molecular Function (MF), and Cellular Component (CC). To retain the complete term set for downstream analysis, enrichment was initially run without internal multiple-testing filtering (pAdjustMethod = “none”, pvalueCutoff = 1, qvalueCutoff = 1) on a defined background. Where reported, Benjamini Hochberg FDR correction was applied post hoc. Enrichment results were visualized with the enrichplot package.

### Correlation between 5xFAD transcriptomics and proteomics

We used log2FC based on the correlation between RNA expression and protein abundance in 5xFAD mice by analyzing the overlapping genes from both datasets using pearson correlation and linear regression.

### Ingenuity Pathway Analysis (IPA) for Transcriptomics and Proteomics

Differentially expressed genes (DEGs) from transcriptomic profiling and differentially expressed proteins (DEPs) from proteomic analysis (PAP-1 vs. Control) were subjected to pathway enrichment using IPA, QIAGEN). Differentially expressed genes (log2FC > 0.25 or < –0.25, adjusted p < 0.05) and differentially expressed proteins (p < 0.05, log2FC thresholds) from PAP-1 versus control were analyzed in IPA using the transcripts detected genes or proteins as background. Canonical Pathways, upstream Regulators, and diseases and functions modules were queried, and activation z-scores were used to infer pathway activity across transcriptomic and proteomic levels.

### snRNA seq derived cell type based Upstream regulators prediction using IPA

We also performed regulatory effect analysis to predict the upstream regulator(s) that may drive functional differences for each cell type in PAP-1 vs. control. The IPA Upstream Regulator module was applied to cell type based pseudobulk differentially expressed genes (log2 fold-change > 0.25 or < –0.25, adjusted p < 0.05) derived from the snRNA-seq Seurat object (PAP-1 vs. control). This analysis predicted transcription factors, cytokines, and signaling molecules responsible for the observed expression changes and inferred their activation or inhibition states based on the directionality of regulated targets.

### Eigen based resilience study in PAP-1 versus Control

We utilized resilience-related protein lists derived from the proteome-wide association study (PWAS) of cognitive trajectory conducted by Yu et al., which examined 8,356 proteins for links with longitudinal cognitive performance. Proteins meeting an FDR threshold of <0.2 were selected, designating those positively associated as pro-resilience (n = 300) and those negatively associated as anti-resilience (n = 216). These sets were intersected with proteins that were differentially up and downregulated in PAP-1 compared to control. From this overlap, 25 pro-resilience proteins upregulated in PAP-1 were used to construct an eigen-based module. These synthetic eigen components were extracted to represent the dominant expression trend within this subset, and eigen scores were compared between PAP-1 and control using a two-tailed t-test.

### Analysis of existing mouse TMT data

Cortical brain tissue from both wild-type and 5xFAD mice underwent a 16-plex tandem mass tag labeling workflow for quantitative proteomic analysis. Proteins were extracted, digested, and labeled before being separated through liquid chromatography and subsequently analyzed by tandem mass spectrometry. The samples were evaluated in six separate batches, each incorporating a pooled global reference sample for normalization across the runs. Quantification at the protein level was carried out for all samples, resulting in the detection of 8,535 proteins. The quantified dataset featured the human Aβ42 peptide, along with proteins linked to amyloid pathology and glial reactions such as GFAP, SV2A, IBA1, in addition to signaling-related proteins like STAT1 and STAT3. Normalized abundance values were utilized for further statistical analysis.

### Other sources of data used for analyses in this manuscript

We used human CSF AD dataset for comparison analysis^61^and Human Post-mortem brain proteomic data were obtained from Johnson et. al.^90^

### Other statistical considerations

Specific statistical tests used for individual experiments are described in the corresponding figure of legends. In general, continuous variables were analyzed using parametric statistical tests, including two-tailed unpaired *t*-tests assuming equal variances for comparisons between two groups, and one-way analysis of variance (ANOVA) followed by Tukey’s honestly significant difference (HSD) post-hoc test for comparisons involving more than two groups. For RNA-sequencing analyses, differential expressions were assessed using DESeq2, with statistical significance determined based on adjusted *p*-values (*padj*) to account for multiple testing.

## Data availability

Brain bulk proteomics data is available on PXD071599(Bulk brain proteomics) PXD072055(CSF proteomics).

## Acknowledgements

We thank Ruth S. Nelson for IHC image analysis.

## Contributions

Conceptualization: SR, RK, CB

Methodology and investigation: RK, US, CB, ADB, PK, DK, EWJ, MB, HZ, SM, AS, SAS

Writing-Original draft: RK, SR, CB, US

Writing-Review and Editing: SR, RK, HW Funding acquisition: SR

Resources: SR, HW

