## Supplementary figure and legends for "Kv1.3 inhibition alleviates neuropathology via neuroinflammatory and resilience pathways in a mouse model of Aβ pathology"

[#]Equally contributed as first author

*Corresponding author

Correspondence to:

Supplemental Figures: 6

Supplemental Datasheets:

Keywords: Alzheimer's disease, Microglia, Kv1.3, Transcriptomics, snRNA seq Proteomics, Interferon, Neuroinflammation, Resilience, CSF Biomarkers

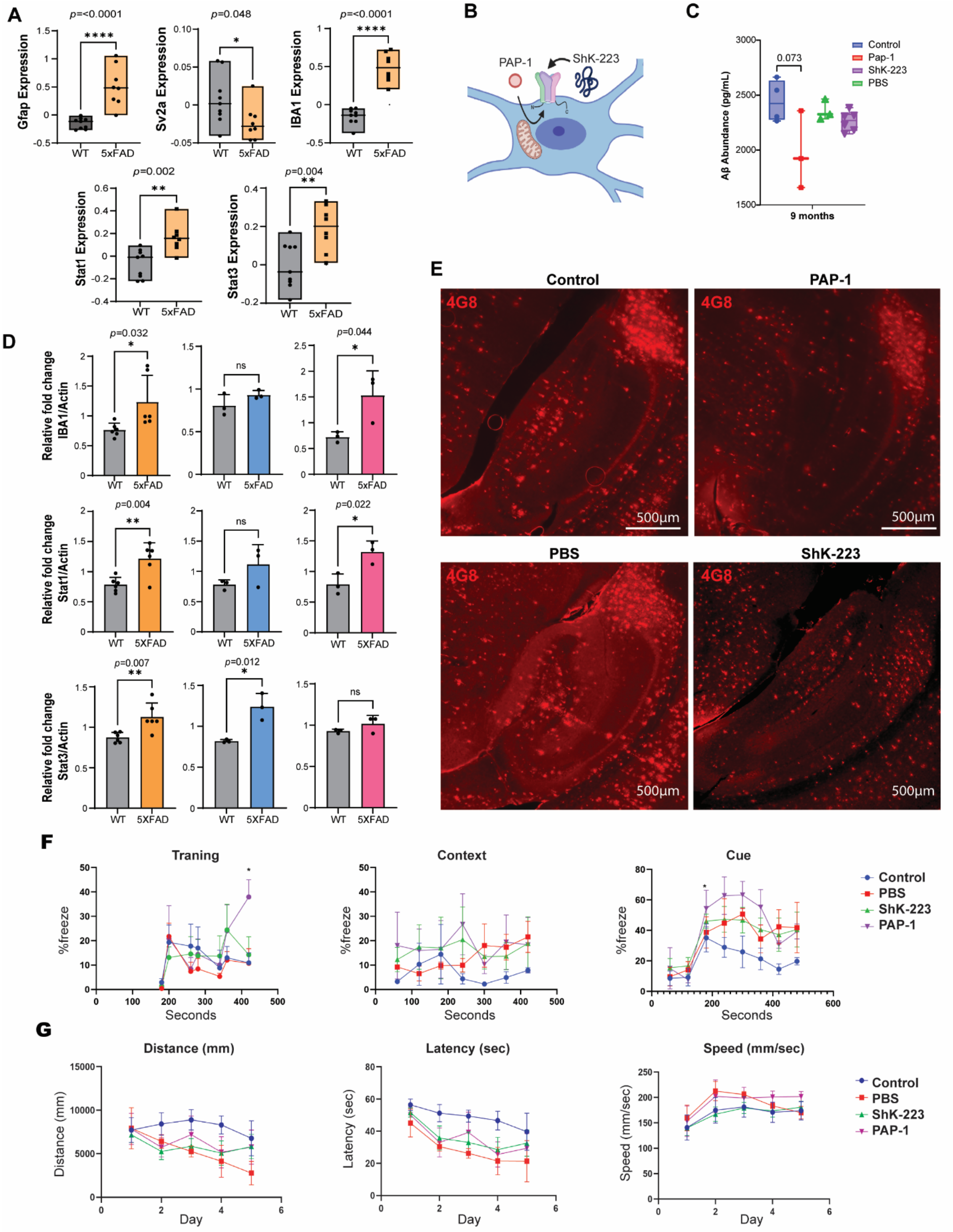

A
p=<0.0001
****
Gfap Expression
WT
5xFAD
p=0.048
*
Sv2a Expression
p=<0.0001
****
IBA1 Expression
p=0.002
**
Stat1 Expression
p=0.004
**
Stat3 Expression
B
PAP-1
ShK-223
C
Aβ Abundance (pg/mL)
0.073
Control
Pap-1
ShK-223
PBS
9 months
D
Relative fold change IBA1/Actin
Relative fold change Stat1/Actin
Relative fold change Stat3/Actin
p=0.032
p=0.044
p=0.004
p=0.022
p=0.007
p=0.012
ns
5XFAD
E
Control
PAP-1
PBS
ShK-223
4G8
500µm
F
Traning
Context
Cue
%freeze
Seconds
Control
PBS
ShK-223
PAP-1
G
Distance (mm)
Latency (sec)
Speed (mm/sec)
Day

**Supplementary Figure 1: Effects of Kv1.3 blockers on neuropathology and behavior (A)** Protein abundance of markers of synaptic integrity, gliosis and neuroinflammation from 10-month-old WT and 5xFAD mice (Male/Female) using TMT/MS **(B)** Biorender diagram of blockade of Kv1.3. PAP-1 crosses the cellular membrane and binds to the inner portion of the Kv1.3 channel while ShK-223 binds to the external region of Kv1.3. Both result in a reduction of potassium flow out of the channel. **(C)** Aβ ELISA shows a nonsignificant trend of reduction of total Aβ with PAP-1. **(D)** Densitometric analysis of western blots in fig1B **(E)** Brain tissue stained for Aβ, (n=3, 3 sections per mouse) and imaged for hippocampal region at 4x for quantification using the Keyence. **(F)** Fear conditioning evaluation of 9-month-old mice shows a difference in the training capacity (association of cue (light) with context (shock)) of PAP-1 treated mice. PAP-1 increases the training capacity of 5xFAD mice. The association of the cue (light) and context (shock) shows no change. **(G)** Morris water maze shows a nonsignificant trend of blockade of Kv1.3, through ShK-223 or PAP-1 decreasing the distance mice traveled before finding the platform and the latency (time present in the water). * $p<0.05$, ** $p < 0.01$, *** $p<0.001$.

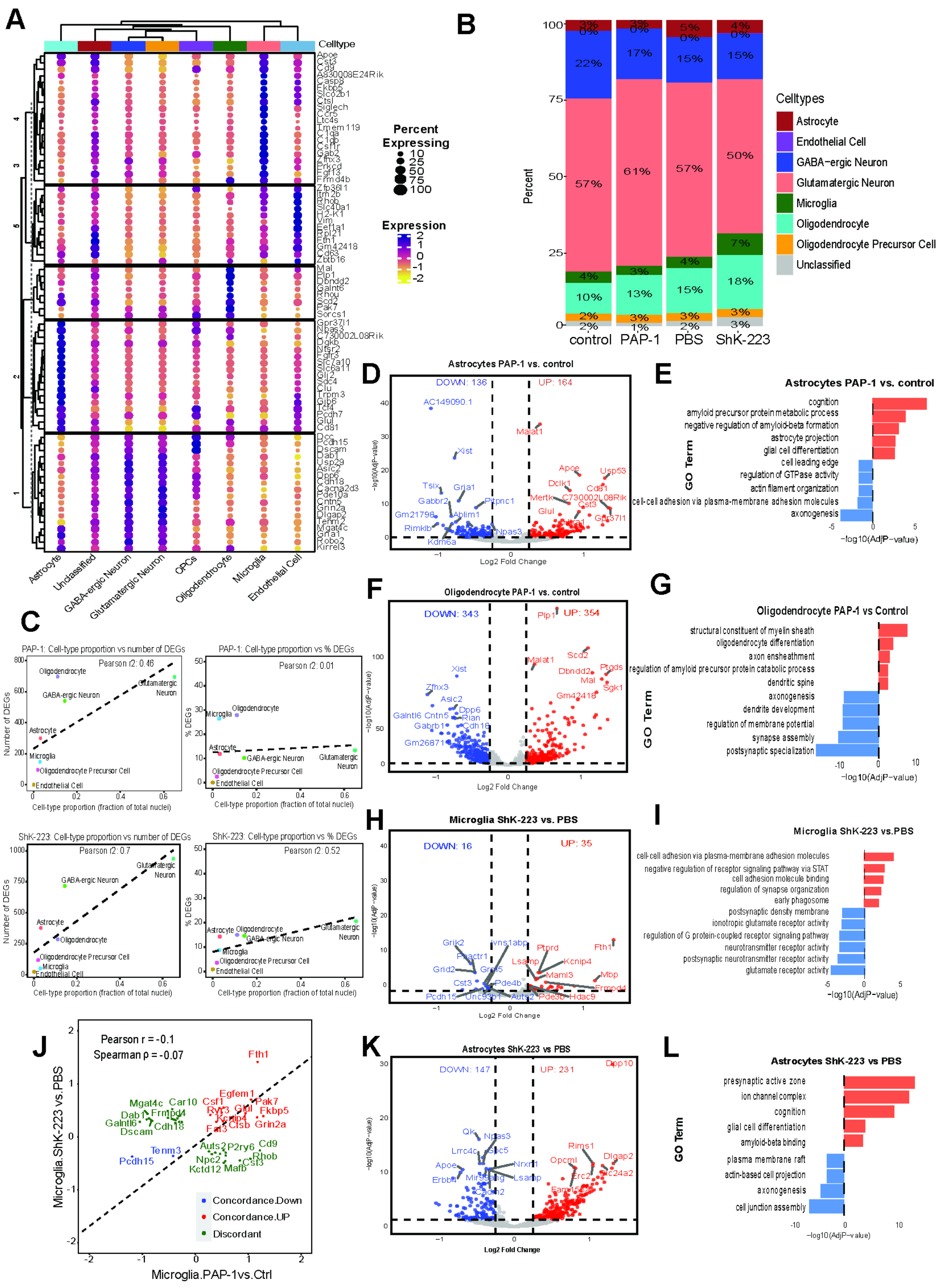

A
Celltype
Percent Expressing
10
25
50
75
100
Expression
2
1
0
-1
-2
Astrocyte
Unclassified
GABA-ergic Neuron
Glutamatergic Neuron
OPCs
Oligodendrocyte
Microglia
Endothelial Cell
B
Percent
control
PAP-1
PBS
ShK-223
Celltypes
Astrocyte
Endothelial Cell
GABA−ergic Neuron
Glutamatergic Neuron
Microglia
Oligodendrocyte
Oligodendrocyte Precursor Cell
Unclassified
C
PAP-1: Cell-type proportion vs number of DEGs
Pearson r2: 0.46
PAP-1: Cell-type proportion vs % DEGs
Pearson r2: 0.01
ShK-223: Cell-type proportion vs number of DEGs
Pearson r2: 0.7
ShK-223: Cell-type proportion vs % DEGs
Pearson r2: 0.52
Number of DEGs
% DEGs
Cell-type proportion (fraction of total nuclei)
D
Astrocytes PAP-1 vs. control
DOWN: 136
UP: 164
Log2 Fold Change
E
Astrocytes PAP-1 vs. control
cognition
amyloid precursor protein metabolic process
negative regulation of amyloid-beta formation
astrocyte projection
glial cell differentiation
cell leading edge
regulation of GTPase activity
actin filament organization
cell-cell adhesion via plasma-membrane adhesion molecules
axonogenesis
GO Term
−log10(AdjP−value)
F
Oligodendrocyte PAP-1 vs. control
DOWN: 343
UP: 354
G
Oligodendrocyte PAP-1 vs Control
structural constituent of myelin sheath
oligodendrocyte differentiation
axon ensheathment
regulation of amyloid precursor protein catabolic process
dendritic spine
axonogenesis
dendrite development
regulation of membrane potential
synapse assembly
postsynaptic specialization
H
Microglia ShK-223 vs. PBS
DOWN: 16
UP: 35
I
Microglia ShK-223 vs.PBS
cell-cell adhesion via plasma-membrane adhesion molecules
negative regulation of receptor signaling pathway via STAT
cell adhesion molecule binding
regulation of synapse organization
early phagosome
postsynaptic density membrane
ionotropic glutamate receptor activity
regulation of G protein-coupled receptor signaling pathway
neurotransmitter receptor activity
postsynaptic neurotransmitter receptor activity
glutamate receptor activity
J
Pearson r = -0.1
Spearman ρ = -0.07
Microglia.ShK-223 vs. PBS
Microglia.PAP-1vs.Ctrl
Concordance.Down
Concordance.UP
Discordant
K
Astrocytes ShK-223 vs PBS
DOWN: 147
UP: 231
L
Astrocytes ShK-223 vs PBS
presynaptic active zone
ion channel complex
cognition
glial cell differentiation
amyloid-beta binding
plasma membrane raft
actin-based cell projection
axonogenesis
cell junction assembly

**Supplementary Figure 2: snRNA seq derived cell type specific pseudo bulk shows transcriptional responses to PAP-1 and ShK-223 (A)** Dot plot of Cell type specific markers expression across major cell populations. Dot size reflects the proportion of nuclei expressing each gene, while color intensity indicates the average expression level within that cluster. **(B)** Stacked bar plot showing the percentage composition of major brain cell types (neurons, glia, vascular cells, OPCs) across control, PAP-1, PBS, and ShK-223 groups. Each colored segment represents the relative abundance of a specific cell type within each condition **(C)** Top panels show PAP-1 cell-type abundance versus transcriptional changes while bottom panels in ShK-223 cell type abundance versus transcriptional response. Scatter plots show the relationship between cell-type proportion (fraction of total nuclei) and transcriptional output following either PAP-1or Shk-223 treatment. The top and bottom left panel displays cell-type proportion versus the absolute number of differentially expressed genes, in both treatment while the top and bottom right panel show cell-type proportion versus the percentage of differentially expressed genes relative to the total genes detected per cell type in both treatments. Each point represents a major brain cell type, and dashed lines indicate linear regression fits. **(D)**Volcano plot of Pseudo-bulk differentially expressed genes in astrocytes comparing PAP-1 with control. Significantly upregulated transcripts are shown in red and downregulated transcripts in blue, with selected genes labeled. The x-axis represents $\log_2$ fold change, and the y-axis represents $\log_{10}$ adjusted p-value, indicating statistical significance of gene expression changes. **(E)** Gene Ontology enrichment of astrocytic transcripts altered by PAP-1 versus Control. Bar length reflects the strength of enrichment ($-\log_{10}$ adjusted P-value) with red color bars representing upregulated pathways and blue bar representing downregulated pathways. **(F)** Volcano plot of pseudo-bulk differentially expressed genes in oligodendrocytes comparing PAP-1 with Control. Significantly upregulated transcripts are shown in red and downregulated transcripts in blue, with selected genes labeled. The x-axis represents $\log_2$ fold change, and the y-axis represents $-\log_{10}$ adjusted p-value, indicating statistical significance of gene expression changes. **(G)** Gene Ontology enrichment of oligodendrocyte transcripts altered by PAP-1 versus Control. Longer bars indicate pathways with stronger enrichment ($-\log_{10}$ adjusted P-value).**(H)** Volcano plot of pseudo-bulk differentially expressed genes in microglia comparing ShK-223 treatment with PBS. Upregulated genes are shown in red and downregulated genes in blue, with selected transcripts labeled. **(I)** Gene Ontology enrichment of microglial transcripts after ShK-223 versus PBS. Longer bars indicate stronger enrichment. **(J)** Concordance analysis of microglial transcripts comparing PAP-1 versus Control and ShK-223 versus PBS. Each point represents a gene, with axes showing $\log_2$ fold change in the two conditions. Genes consistently upregulated, consistently downregulated, showing opposite regulations are highlighted. **(K)** Volcano plot of pseudo-bulk differentially expressed genes in astrocytes comparing ShK-223 with PBS. Upregulated genes are shown in red and downregulated genes in blue, with selected transcripts labeled. The x-axis shows $\log_2$ fold change, and the y-axis shows $-\log_{10}$ adjusted p-value. **(L)** GO enrichment in astrocytes treated with ShK-223 compared with PBS.

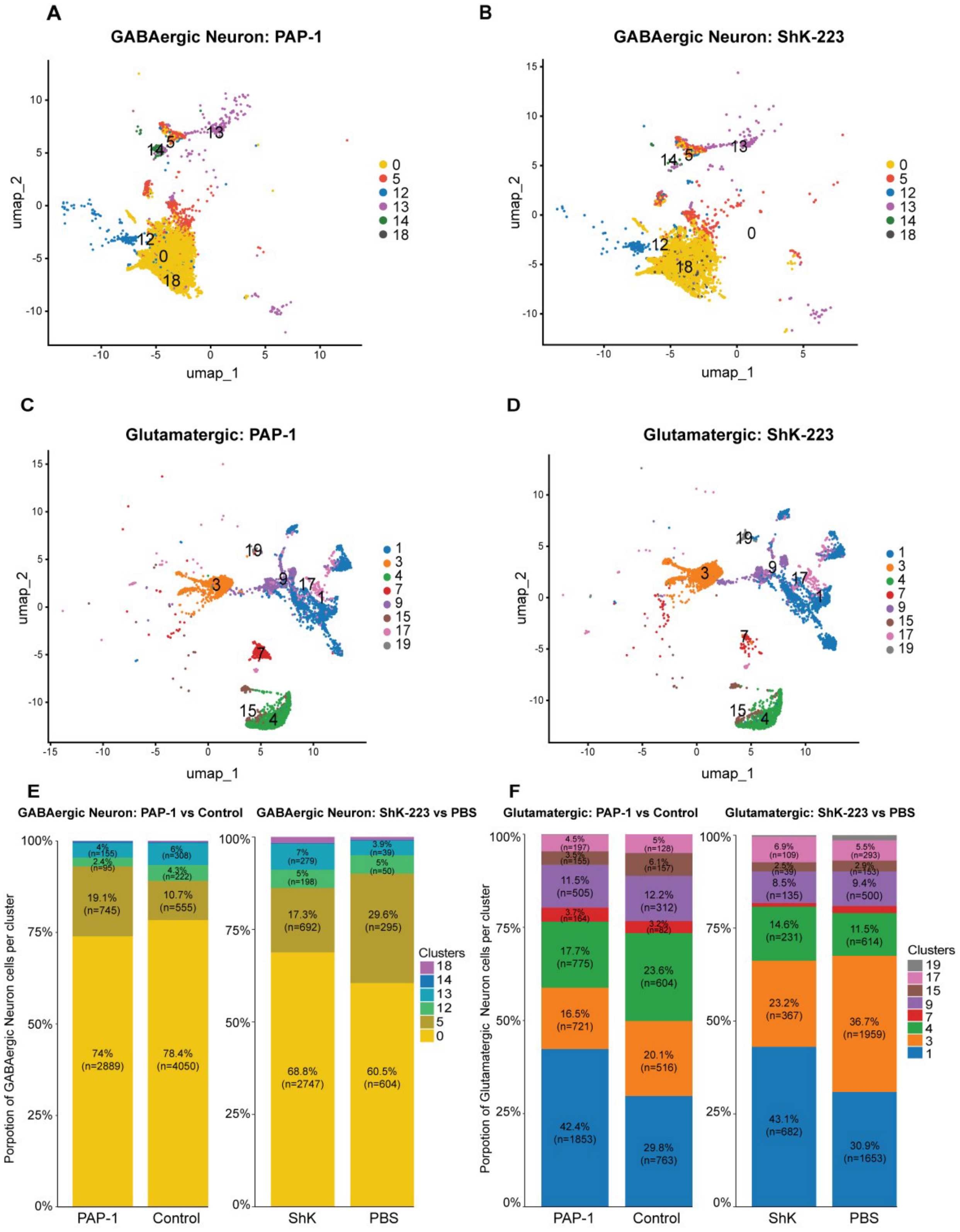


| | Celltype | Chi-Square | P-value |
|---|---|---|---|
| PAP-1 vs Control | GABAergic | 157.5 | 2.2e-16 |
| ShK-223 vs PBS | GABAergic | 84.28 | 2.2e-16 |

| | Celltype | Chi-Square | P-value |
|---|---|---|---|
| PAP-1 vs Control | Glutamatergic | 135.61 | 2.2e-16 |
| ShK-223 vs PBS | Glutamatergic | 151.53 | 2.2e-16 |

**Supplementary Figure 3: Effects of PAP-1 and ShK-223 on GABAergic and glutamatergic neuronal cluster composition (A)** UMAP projection of GABAergic neurons under PAP-1 treatment, colored by cluster identity (clusters 0, 5, 12, 13, 14, and 18). Distinct inhibitory neuronal clusters are preserved, indicating maintained GABAergic identity, with treatment-associated differences in cluster distribution. **(B)** UMAP visualization of GABAergic neurons following ShK-223 treatment, showing comparable cluster identities to PAP-1 treated neurons. Changes in cluster localization and relative cluster distribution suggest differential modulation of inhibitory neuronal subpopulations by ShK-223treatment**. (C)** Glutamatergic neuronal subclusters in the PAP1 condition are illustrated in UMAP, where each point indicates a single cell colored by cluster detection. Within excitatory neuronal populations, distinct clusters (1, 3, 4, 7, 9, 15, 17, and 19) show PAP1-associated remodeling and transcriptional change. **(D)** UMAP visualization of glutamatergic neurons following ShK-223 treatment, colored by cluster identity (clusters 1, 3, 4, 7, 9, 15, 17, and 19). Redistribution of excitatory neuronal clusters is observed while overall cluster structure is preserved, indicating treatment-dependent modulation of glutamatergic neuronal subpopulations without loss of excitatory identity. **(E)** The relative proportions of GABAergic neuron clusters are displayed in stacked bar charts comparing PAP-1 vs. Control (left) and ShK-223 vs. PBS (right). Chi-square analysis indicates that these differences are statistically significant (PAP-1 vs. Control: $\chi^2 = 157.5$, $p < 2.2 \times 10^{-1\circ}$; ShK-223 vs. PBS: $\chi^2 = 84.28$, $p < 2.2 \times 10^{-11}$). **(F)** The relative abundance of glutamatergic neuron clusters under PAP-1 vs. Control (left) and ShK-223 vs. PBS (right) conditions is shown in stacked bar plots. For both comparisons (PAP-1 vs. Control: $\chi^2 = 135.61$, $p < 2.2 \times 10^{-1\circ}$; ShK-223 vs. PBS: $\chi^2 = 151.53$, $p < 2.2 \times 10^{-11}$), chi-square testing highlights a significant distribution of cluster composition.

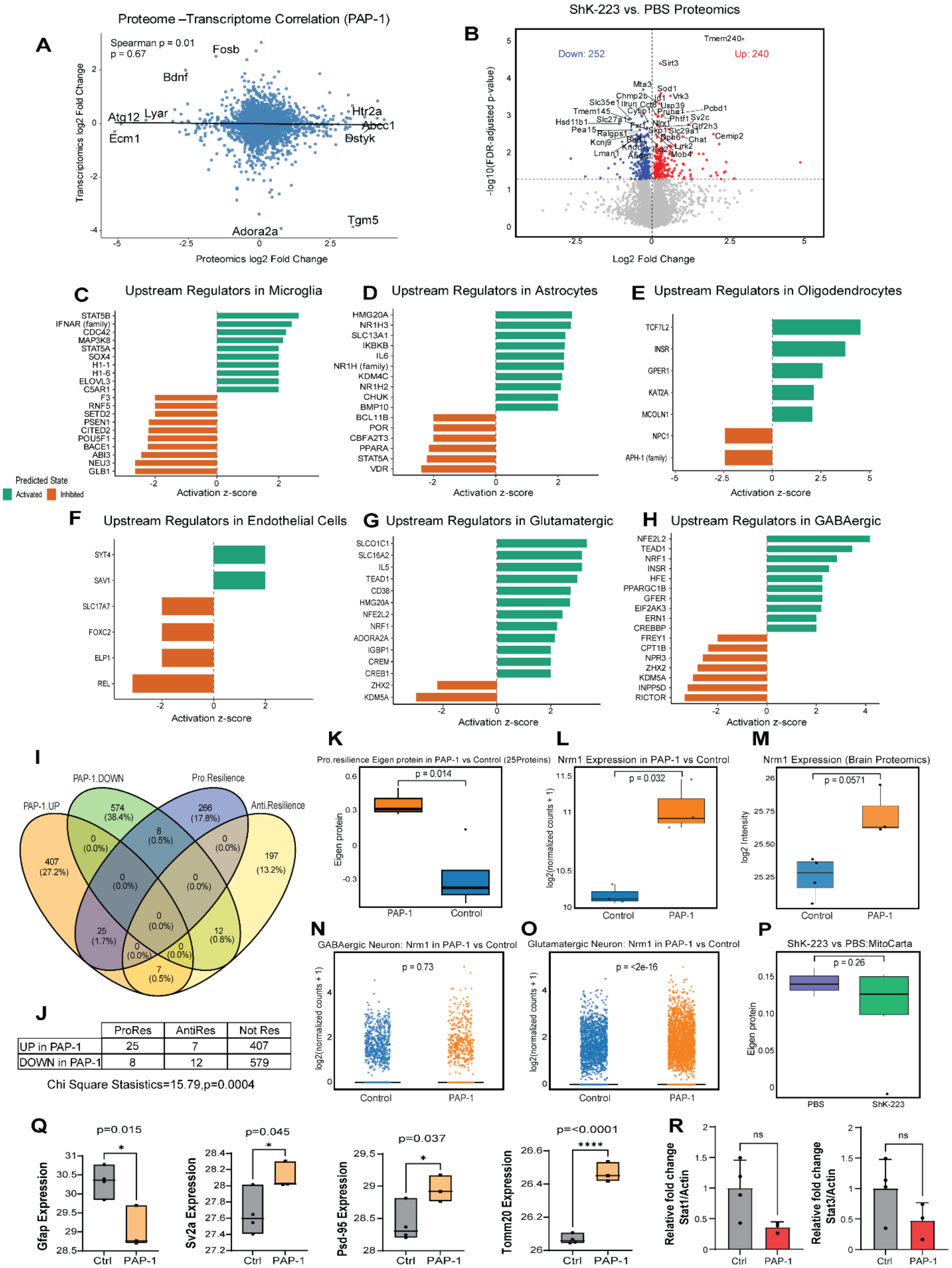


| | ProRes | AntiRes | Not Res |
|---|---|---|---|
| UP in PAP-1 | 25 | 7 | 407 |
| DOWN in PAP-1 | 8 | 12 | 579 |

Chi Square Stasistics=15.79,p=0.0004

**Supplementary Figure 4: Cell type specific upstream regulation analyses and resilience-linked mechanisms in PAP-1 (A)** Transcriptomic and proteomic datasets from the brains of PAP-1-treated and control mice were correlated. Overall consistency in mRNA and protein abundance levels is shown by each point, which represents a gene–protein pair. **(B)** Volcano plot showing differentially expressed proteins in the brain proteomics following ShK-223 treatment compared with PBS, with 252 proteins downregulated and 240 upregulated. (C-H) Cell-type specific upstream regulators predicted by snRNA seq for the main types of brain cells. Bar plot showing predicted upstream regulators active in endothelial transcriptomic signatures. Y-axis lists upstream regulators, and X-axis represents activation of z-score. Green bars indicate predicted activation (positive z-score), whereas orange bars represent predicted inhibition (negative z-score) (**C)** Upstream regulators in microglia. **(D)** Upstream regulators in Astrocytes. **(E)** Upstream regulators in Oligodendrocytes. **(F)** Upstream regulators in Endothelial cells**. (G)** Upstream regulators in Glutamatergic neurons. **(H)** Upstream regulators in GABAergic Neurons **(I)** Resilience-associated modules overlap with proteins controlled by PAP-1. Venn diagram illustrates the intersection of pro-, anti-, and PAP-1-downregulated (PAP-1 DOWN) and PAP-1-upregulated (PAP-1 UP) protein modules obtained from the proteome dataset. While there is barely any overlap with anti-resilience proteins, eight proteins show agreement between PAP-1 DOWN and pro-resilience modules, indicating that PAP-1 treatment alters molecular networks leading to increased resilience. **(J)** The distribution of pro-resilience, anti-resilience proteins in PAP-1 upregulated and downregulated proteins summarized in table with chi-square test ($\chi^2$ = 15.79, p = 0.0004).**(K)** Boxplot showing higher pro-resilience eigen protein values in PAP-1 compared with control. **(L-M)** Validation of Nrn1 expression following PAP-1 treatment. Left: Transcriptomic analysis revealed a significant increase in *Nrn1* transcript levels in PAP-1–treated samples compared to control ($p$ = 0.032). Right: Proteomic analysis shows a similar upward trend in NRN1 protein abundance ($p$ = 0.0571), supporting transcriptional upregulation of Nrn1 and its potential role in PAP-1–induced resilience enhancement. (**N)** Boxplot showing no significant change in GABAergic neuron *Nrn1* expression between PAP-1 and control. **(O)** Boxplot showing significantly higher *Nrn1* expression in glutamatergic neurons with PAP-1 treatment. **(P)** Boxplot showing no reduction in MitoCarta eigen protein values in ShK-223 versus PBS. **(Q)** Boxplots of GFAP, SV2A, PSD-95 and TOMM20 in Control vs. PAP-1from bulk proteomics MS data. The y-axis shows normalized expressions. Boxes mark median and interquartile range; whiskers show replicate spread. PAP-1 samples show significant upregulation (**R)** Densitometric analysis of STAT1 and STAT3 from fig 5K.

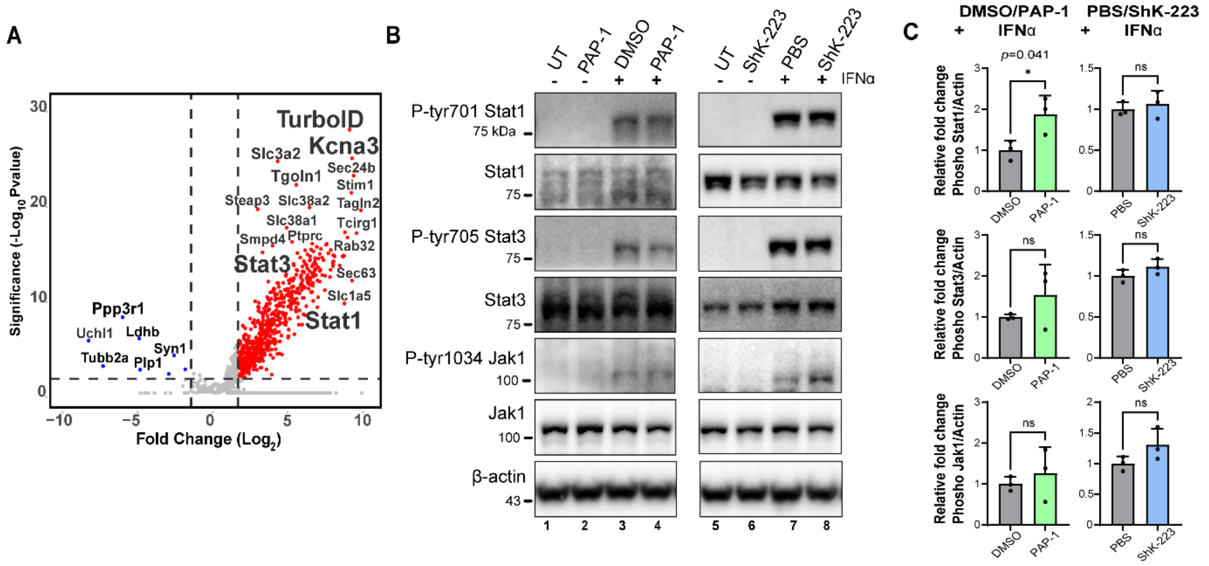


**Supplementary Figure 5: Identification of Kv1.3–STAT interactions and assessment of PAP-1/ShK-223 effects on IFNα-mediated STAT/JAK phosphorylation (A)** Volcano plot showing TurboID-enriched proteins several protein-protein interactors of Kv1.3 in microglia were identified, including STAT proteins which are key regulators in interferon signaling from Bowen *et al* dataset[1] **(B, C)** Western blot and densitometric analysis of whole cell lysate of interferon α induced MMC cells post Kv1.3 blockade by PAP-1 or ShK-223 shows no effect on phosphorylation of STAT1, STAT3 and JAK1.

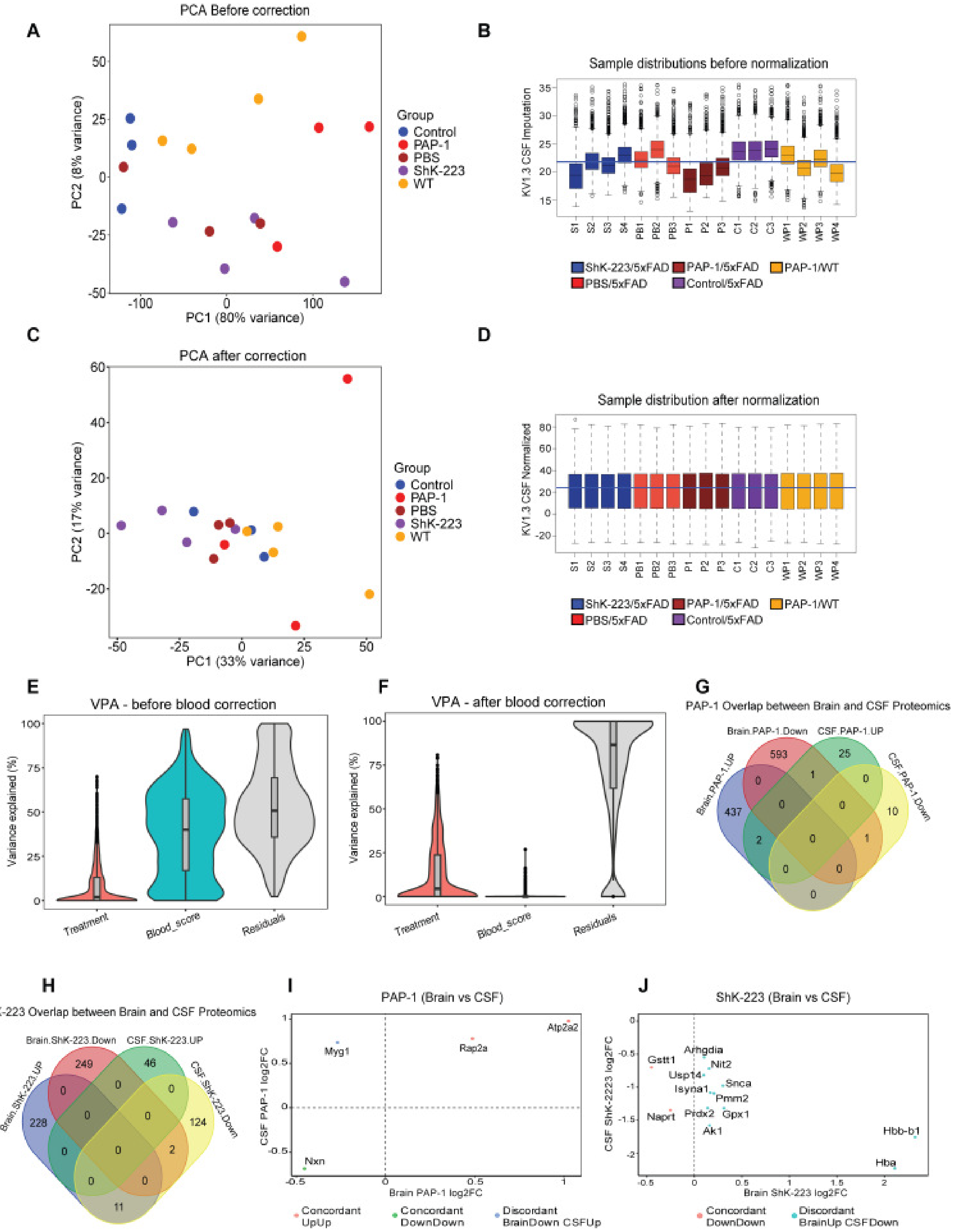

A
PCA Before correction
PC1 (80% variance)
PC2 (8% variance)
Group
Control
PAP-1
PBS
ShK-223
WT
B
Sample distributions before normalization
KV1.3 CSF Imputation
S1 S2 S3 S4 PB1 PB2 PB3 P1 P2 P3 C1 C2 C3 WP1 WP2 WP3 WP4
ShK-223/5xFAD
PAP-1/5xFAD
PAP-1/WT
PBS/5xFAD
Control/5xFAD
C
PCA after correction
PC1 (33% variance)
PC2 (17% variance)
Group
Control
PAP-1
PBS
ShK-223
WT
D
Sample distribution after normalization
KV1.3 CSF Normalized
S1 S2 S3 S4 PB1 PB2 PB3 P1 P2 P3 C1 C2 C3 WP1 WP2 WP3 WP4
ShK-223/5xFAD
PAP-1/5xFAD
PAP-1/WT
PBS/5xFAD
Control/5xFAD
E
VPA - before blood correction
Variance explained (%)
Treatment
Blood_score
Residuals
F
VPA - after blood correction
Variance explained (%)
Treatment
Blood_score
Residuals
G
PAP-1 Overlap between Brain and CSF Proteomics
Brain.PAP-1.Down
CSF.PAP-1.UP
Brain.PAP-1.UP
CSF.PAP-1.Down
H
ShK-223 Overlap between Brain and CSF Proteomics
Brain.ShK-223.Down
CSF.ShK-223.UP
Brain.ShK-223.UP
CSF.ShK-223.Down
I
PAP-1 (Brain vs CSF)
Brain PAP-1 log2FC
CSF PAP-1 log2FC
Atp2a2
Rap2a
Myg1
Nxn
Concordant UpUp
Concordant DownDown
Discordant BrainDown CSFUp
J
ShK-223 (Brain vs CSF)
Brain ShK-223 log2FC
CSF ShK-2223 log2FC
Gstt1
Arhgdia
Nit2
Usp14
Snca
Isyna1
Pmm2
Naprt
Prdx2
Gpx1
Ak1
Hbb-b1
Hba
Concordant DownDown
Discordant BrainUp CSFDown

**Supplementary Figure 6: CSF Proteomic overview: PCA Shifts, Distribution Normalization, and Removal of Blood-Derived Variation (A)** PCA of the CSF proteome before blood contamination removal shows broad sample dispersion with limited clustering by treatment. PC1 accounts for 80% of the total variation, while PC2 explains 8%, indicating that most of the variation in the raw data is dominated by confounding factors. **(B)** Boxplots of the raw intensity distributions show substantial differences across samples, reinforcing the need for normalization before downstream comparisons. **(C)** After blood contamination removal, the PCA structure changes show PC1 as 33% and PC2 explains 17% of the variation. The shift in explained variance reflects removal of major confounders and a more balanced contribution of biological signals. **(D)** Normalization produces consistent intensity ranges across samples, confirming that global intensity differences were effectively reduced. **(E)** Variance partitioning before blood correction shows that blood-related signal contributes to a large proportion of variability, while treatment effects account for only a small fraction of the total variance**. (F)** After blood correction, the proportion of variance attributed to blood signals is eliminated. Treatment-related variance becomes more interpretable, and the residual component reflects remaining biological and technical noise**. (G)**Venn intersection shows PAP-1 treatment resulted in a large number of brain specific proteomic changes (437 brain upregulated and 593 brain downregulated proteins), whereas CSF changes were limited (25 CSF upregulated and 10 CSF downregulated proteins). Overlap between the brain and CSF was minimal, with only 1 shared protein. **(H)**Venn diagram showing the overlap between differentially expressed proteins in brain and CSF following ShK-223 treatment. Brain proteomics identified 228 upregulated and 249 downregulated proteins, whereas CSF proteomics identified 46 upregulated and 124 downregulated proteins. **(I)** Log2FC Based Scatter plot between brain and CSF responses to PAP-1. Only a small set of proteins was shared across the brain and CSF after PAP-1 treatment, with ATP2A2 and RAP2A increasing in both tissues and NXN decreasing in both, while MYG1 shifted in opposite directions. These patterns produced a positive correlation ($r = 0.723$; $R^2 = 0.522$). **(J)**Log2FC Based Scatter plot between brain and CSF responses to ShK-223. The x-axis represents the log2 fold-change of each protein in the brain after ShK-223 treatment, showing whether its levels increased or decreased. The y-axis shows the $\log_2$ fold-change of the same proteins in the CSF, allowing direct comparison across compartments.


1. Bowen CA*, et al.* Proximity Labeling Proteomics Reveals Kv1.3 Potassium Channel Immune Interactors in Microglia. *Mol Cell Proteomics* **23**, 100809 (2024).
