## Supplementary material for "Kv1.3 inhibition alleviates neuropathology via neuroinflammatory and resilience pathways in a mouse model of Aβ pathology": table legends for tables and supp. tables

### Table Legends: Datasheet 1-7

#### Datasheet 1: Data analysis for Figure 1

**Sheet #1 (Fig.1A):**Tandem mass tag (TMT)-based quantitative proteomics for levels of hAβ peptide levels in WT and 5xFAD brains (10 months).
**Sheet #2 (Fig.1B):**Western blots and quantification of expression of several proteins in whole-brain lysates from 10-month-old WT and 5xFAD mice.
**Sheet #3 (Fig.1C):** Statistical analysis used in the graphs for expressional levels of different protein in WT and 5xFAD mice along with sex specific differences.
**Sheet #4 (Fig.1E):** Quantification of %Aβ coverage in cortical and hippocampal section IHC images of 5xFAD mice in control and PAP1 treatment.
**Sheet #5 (Fig.1F):**Quantification of %Aβ coverage in cortical and hippocampal section IHC images of 5xFAD mice in PBS control and Shk-223 treatment
.

#### Datasheet 2: Data analysis for Figure 2

**Sheet #1 (Fig.2A):** Uniform Manifold Approximation and Projection (UMAP) of integrated single nucleus RNA sequencing displaying the populations of brain cells with annotations based on the expression of canonical marker genes.
**Sheet #2 (Fig.2B):** A summary of the Pseudo-bulk based differentially expressed genes (DEGs) found in microglia after PAP-1 treatment in comparison to the control, highlighting the overall size and direction of transcriptional changes.
**Sheet #3 (Fig.2C):** A summary of microglial Pseudo-bulk DEGs following ShK-223 treatment in comparison to PBS, which enabled transcriptional responses across pharmacologically different Kv1.3 inhibitors to be evaluated.
**Sheet #4 (Fig.2D):** Volcano plot of microglial DEGs in PAP-1 versus control, showing statistically significant genes that are strongly elevated and downregulated together with log2 fold change.
**Sheet #5 (Fig.2E):** Gene ontology (GO) enrichment analysis of microglial DEGs modulated by PAP-1, demonstrating molecular activities and biological processes associated to changes in microglial transcriptional states
**Sheet #6 (Fig.2F):** Venn diagram demonstrating genes that were significantly increased in microglia after PAP-1 treatment in comparison to control.
**Sheet #7 (Fig.2G):** Venn diagram showing genes which were significantly downregulated in microglia after PAP-1 treatment in comparison to the control.
**Sheet #8 (Fig.2H):** Volcano plot of oligodendrocyte DEGs after treatment with ShK-223 versus PBS, demonstrating transcriptional alterations distinctive to this glial lineage
**Sheet #9 (Fig.2I):** GO enrichment analysis of oligodendrocyte DEGs regulated by ShK-223, highlighting pathways associated with myelin-related and oligodendrocyte biology.
**Sheet #10 (Fig.2J):** Venn diagram showing genes significantly increased in oligodendrocytes after treatment with ShK-223.
**Sheet #11 (Fig.2K):** Venn diagram shows genes significantly downregulated in oligodendrocytes after treatment with ShK-223.

#### Datasheet 3: Data analysis for Figure 3

**Sheet #1 (Fig.3A):** Analysis of glutamatergic neurons' pseudo bulk differential expressions comparing PAP-1 and control samples.
**Sheet #2 (Fig.3B):** PAP-1 regulated transcripts in glutamatergic neurons are enriched in gene ontology.
**Sheet #3 (Fig.3C):** Glutamatergic and GABAergic neurons share PAP-1-regulated differentially expressed genes.
**Sheet #4 (Fig.3D):** GABAergic neurons pseudo bulk differential expression analysis comparing PAP-1 and control samples.

**Sheet #5 (Fig.3E):** PAP-1-regulated transcripts in GABAergic neurons are enriched in gene ontology.
**Sheet #6 (Fig.3F):** Correlation between differentially expressed genes in GABAergic neurons controlled by PAP-1 and ShK-223.
**Sheet #7 (Fig.3G):** ShK-223 and PBS samples were compared in a pseudo bulk differential expression study involving glutamatergic neurons
**Sheet #8 (Fig.3H):** ShK-223-regulated transcripts in glutamatergic neurons are enriched in gene ontology.
**Sheet #9 (Fig.3I):** ShK-223-regulated differentially expressed genes in glutamatergic and GABAergic neurons overlap.
**Sheet #10 (Fig.3J):** GABAergic neurons' pseudo bulk differential expression analysis comparing ShK-223 and PBS samples.
**Sheet #11 (Fig.3K):** ShK-223-regulated transcripts in GABAergic neurons are enriched in gene ontology.
**Sheet #12 (Fig.3L):** Correlation table comparing log2 fold changes of PAP-1 and ShK-223 regulated differentially expressed genes in glutamatergic neurons, assessing concordance and discordance between Kv1.3 inhibitors

### Datasheet 4: Data analysis for Figure 4

**Sheet #1 (Fig.4A):** Cell-cell interaction of ligand and receptors across source and targets inferred following PAP-1 treatment, depicting changes in the number and directionality of ligand and receptor interactions among major brain cell types.
**Sheet #2 (Fig.4B):** Endothelial to neuron ligand receptor (LR) interactions following PAP-1 treatment.
**Sheet #3 (Fig.4C):** Cell-cell interaction ligand and receptors across source and targets inferred following ShK-223 treatment compared with PBS, illustrating treatment associated changes in intercellular communication across major brain cell types.
**Sheet #4 (Fig.4D):** Ligand-receptors across endothelial neuron cell papulation in response to ShK-223 Treatment.
**Sheet #5(Fig.4E):** Heatmap of significantly upregulated ligand-receptor pairs in PAP-1 treated samples compared with control (P value < 0.05), showing source to target cell type interactions and interaction probabilities.
**Sheet #6 (Fig.4F):** Heatmap of significantly downregulated ligand-receptor pairs in PAP-1 treated samples compared with control (P value < 0.05).
**Sheet #7 (Fig.4G):** Heatmap of significantly upregulated ligand receptor pairs in ShK-223 treated samples compared with PBS (P < 0.05).
**Sheet #8 (Fig.4H):** Heatmap of significantly downregulated ligand-receptor pairs in ShK-223 treated samples compared with PBS (P < 0.05).

### Datasheet 5: Data analysis for Figure 5

**Sheet #1 (Fig.5A):** Bulk brain transcriptomic differential expression analysis comparing PAP-1–treated control samples.
**Sheet #2 (Fig.5B):** Gene ontology enrichment analysis of upregulated and downregulated genes identified in bulk brain transcriptomics following PAP-1 treatment.
**Sheet #3 (Fig.5C):** Ingenuity Pathway Analysis of transcriptomic changes associated with PAP-1 treatment.
**Sheet #4 (Fig.5D):** Differentially expressed proteins identified by quantitative proteomics comparing PAP-1 and control samples
**Sheet #5 (Fig.5E):** Gene ontology enrichment analysis of PAP-1–regulated proteins.
**Sheet #6 (Fig.5F):** Ingenuity Pathway Analysis of proteomic changes associated with PAP-1 treatment.
**Sheet #7 (Fig.5G):** Protein list corresponding to synaptic membrane associated proteins upregulated by PAP-1

**Sheet #8 (Fig.5H):** Protein list corresponding to transcription factor binding–associated proteins downregulated by PAP-1.
**Sheet #9 (Fig.5I):** Heatmap matrix of the top 25 resilience-associated proteins comparing control and PAP-1 samples
**Sheet #10 (Fig.5J):** Boxplot of eigenprotein expression for Mito Carta-associated proteins following PAP-1 treatment.
**Sheet #11(Fig.5K):** Quantification of control and PAP-1 treated mice for expression of different proteins of synaptic integrity, gliosis, mitochondria, and inflammation
**Sheet #12(Fig.5L):** Statistical analysis of western blot quantification of different proteins of Fig 5K

### Datasheet 6: Data analysis for Figure 6

**Sheet #1 (Fig.6A):** Western blots showing co immunoprecipitation of Kv1.3 and STATs
**Sheet #2 (Fig.6B):** Western blot and quantification of cell lysate of BV2 SR73 upon Kv1.3 blockade using PAP-1 or Shk-223 followed by IFNγ induction.
**Sheet #3 (Fig.6C):** Statistical analysis used in the graphs for expressional levels of different proteins from figure 6B
**Sheet #4 (Fig.6D):** Western blot and quantification of cell lysate of MMC upon Kv1.3 blockade using PAP-1 or Shk-223 followed by IFNγ induction
**Sheet #5 (Fig.6E):** Statistical analysis used in the graphs for expressional levels of different proteins from figure 6D
**Sheet #6 (Fig.6F)** Western blot and quantification of cell lysate of BV2 SR73 upon Kv1.3 blockade using PAP-1 or Shk-223 followed by IFNα induction.
**Sheet #7 (Fig.6G)** Statistical analysis used in the graphs for expressional levels of different proteins from figure 6D
**Sheet #8 (Fig.6H)** Western blot and quantification of cell lysate of WT and *Kcna3* KO IFN γ induction
**Sheet #9 (Fig.6I)** Statistical analysis used in the graphs for expressional levels of different proteins from figure 6H

### Datasheet 7: Data analysis for Figure 7

**Sheet #1 (Fig.7A)** CSF differentially expressed proteins (DEPs) between 5xFAD and WT mice.
**Sheet #2 (Fig.7B)** CSF differentially expressed proteins (DEPs) between PAP1-treated and control mice.
**Sheet #3 (Fig.7C)** CSF proteomics analysis identifying differentially expressed proteins (DEPs) between ShK-223-treated and PBS-treated mice.
**Sheet #4 (Fig.7D)** Gene Ontology (GO) enrichment analysis of CSF proteins altered in 5xFAD mice relative to wild type.
**Sheet #5 (Fig.7E)** Gene Ontology (GO) enrichment analysis of PAP-1 regulated CSF proteins relative to control.
**Sheet #6 (Fig.7F)** Gene Ontology (GO) enrichment analysis of ShK-223-regulated CSF proteins relative to PBS.
**Sheet #7 (Fig.7G)** UpSetR Interaction table of overlapping CSF protein signatures across PAP-1 treatment, ShK-223 treatment, and 5xFAD pathology.
**Sheet #8 (Fig.7H)** Cross-species correlation heatmap comparing CSF proteomic alterations in human Alzheimer's disease and mouse PAP1- and ShK-223-treated models, with human CSF data derived from Lenora Higginbotham et al. (2020).

### Supplementary Tables: Supp. Datasheet 1-6

#### Supp. Datasheet 1: Supp.Figure.1 analysis

**Sheet #1 (Suppl.Fig.1A):** Tandem mass tag (TMT)-based proteomics for different markers of gliosis, synaptic integrity and inflammation in WT and 5xFAD brains (10 months)
**Sheet #2 (Suppl.Fig.1C):** Aβ levels in 9 months old mice post PAP-1 and Shk-223 treatment measured using ELISA.
**Sheet #3 (Suppl.Fig.1D):** Statistical analysis of western blots of remaining blots from **Fig1B** that compares WT and 5xFAD mice.
**Sheet #4 (Suppl.Fig.1F):** Raw data table for fear conditioning behavioral test
**Sheet #5 (Suppl.Fig.1G):** Raw data table for Morris water maze behavioral test.

#### Supp. Datasheet 2: Supp.Figure.2 analysis

**Sheet #1 (Suppl.Fig.2B):** Based on integrated single nucleus RNA-sequencing data, relative proportions of major brain cell types throughout experimental settings show similar cell-type representation across treatment groups.
**Sheet #2 (Suppl.Fig. 2C):** The number and percentage of DEGs per cell type plotted against matching cell-type proportions illustrates the relationship between cell-type abundance and transcriptional responsiveness, emphasizing variations in DEG load independent of cell abundance
**Sheet #3 (Suppl.Fig.2D):** A volcano plot of astrocyte DEGs after PAP-1 treatment compared to control, showing $\log_2$ fold change vs statistical significance and highlighting genes that are significantly regulated.
**Sheet #4 (Suppl.Fig.2E):** Gene ontology (GO) enrichment analysis of astrocyte DEGs controlled by PAP-1, showing the molecular activities and biological processes connected to astrocyte transcriptional alterations.
**Sheet #5 (Suppl.Fig.2F):** Volcano plot of oligodendrocyte DEGs after PAP-1 treatment in comparison to control, showing transcriptional alterations related to treatment within the oligodendrocyte lineage.
**Sheet #6 (Suppl.Fig.2G):** GO enrichment analysis of oligodendrocyte DEGs regulated by PAP-1, highlighting pathways related to myelin-associated activities and oligodendrocyte function.
**Sheet #7 (Suppl.Fig.2H):** Volcano plot of microglial DEGs after ShK-223 treatment in comparison to PBS, indicating the extent and importance of microglia transcriptional alterations.
**Sheet #8 (Suppl.Fig.2I):** GO enrichment analysis of microglial DEGs regulated by ShK-223, identifying molecular pathways and biological processes linked to the transcriptional remodeling of microglia in response to ShK-223.
**Sheet #9 (Suppl.Fig.2J):** Correlation analysis comparing the concordance and divergence of transcriptional responses across different Kv1.3 inhibitors by comparing microglial $\log_2$ fold changes induced by PAP-1 and ShK-223.
**Sheet #10 (Suppl.Fig.2K):** A volcano plot of astrocyte DEGs after ShK-223 treatment in comparison to PBS, showing transcriptional alterations unique to astrocytes.
**Sheet #11 (Suppl.Fig.2L):** GO enrichment analysis of astrocyte DEGs controlled by ShK-223 revealed biological mechanisms associated with astrocyte reactions to ShK-223 treatment.

#### Supp. Datasheet 3: Supp.Figure.3 analysis

**Sheet #1 (Suppl. Fig.3A):** GABAergic Neuron cells per cluster in PAP-1 and Control.
**Sheet #2 (Suppl. Fig.3B):** GABAergic Neuron cells per cluster in ShK-223 and PBS.
**Sheet #3 (Suppl. Fig.3C):** Glutamatergic Neuron cells per cluster in PAP-1 and Control.
**Sheet #4 (Suppl. Fig.3D):** Glutamatergic Neuron cells per cluster in ShK-223 and PBS.
**Sheet #5 (Suppl. Fig.3E):** Percentages of GABAergic neuron cells per cluster affected by PAP-1 and Shk-223. Predicted Chi-square values for PAP-1 (X-squared = 84.272) and ShK-223 (X-squared = 157.5).
**Sheet #6 (Suppl. Fig.3F):** Percentages of Glutamatergic neuron cells affected by PAP-1 and Shk-223. Predicted Chi-square values for PAP-1 (X-squared = 135.61) and ShK-223 (X-squared = 151.53).

### Supp. Datasheet 4: Supp.Figure.4 analysis

**Sheet #1 (Suppl.Fig.4A):** Correlation between proteomic and transcriptomic $\log_2$ fold changes following PAP-1 treatment shows minimal concordance, as assessed by Spearman's rank correlation ($\rho = 0.0052$, $p = 0.673$).
**Sheet #2 (Suppl.Fig.4B):** Differentially expressed proteins (DEPs) in brain proteomics comparing ShK-223 and PBS-treated samples.
**Sheet #3 (Suppl.Fig.4C):** Predicted upstream regulators in microglia based on pseudo bulk single-nucleus RNA-sequencing analysis.
**Sheet #4 (Suppl.Fig.4D):** Predicted upstream regulators in astrocytes based on pseudo bulk single-nucleus RNA-sequencing analysis.
**Sheet #5 (Suppl.Fig.4E):** Predicted upstream regulators in oligodendrocytes based on pseudo bulk single-nucleus RNA-sequencing analysis.
**Sheet #6 (Suppl.Fig.4F):** Predicted upstream regulators in endothelial cells based on pseudo bulk single-nucleus RNA-sequencing analysis.
**Sheet #7 (Suppl.Fig.4G):** predicted upstream regulators in glutamatergic neurons based on pseudo bulk single-nucleus RNA-sequencing analysis.
**Sheet #8 (Suppl.Fig.4H):** Predicted upstream regulators in GABAergic neurons based on pseudo bulk single-nucleus RNA-sequencing analysis.
**Sheet #9(Suppl.Fig.4I):** Four-way Venn analysis showing overlap of PAP-1 regulated genes with pro-resilience and anti-resilience gene sets.
**Sheet#10(Suppl.Fig.4J):** A Chi-square test revealed a significant association between PAP-1–induced regulation (UP vs DOWN) and resilience category (Pro-resilience, Anti-resilience, Not-resilient) ($\chi2 = 12.02$, $p = 0.00245$).
**Sheet#11(Suppl.Fig.4K):** Pro-resilience eigen-protein expression comparing PAP-1 and control samples, derived from a 25-protein signature.
**Sheet#12(Suppl.Fig.4L)** Nrn1 expression in bulk transcriptomic data from 5xFAD mice comparing control and PAP-1 treatment shows P-value =0.014.
**Sheet#13(Suppl.Fig.4M)** Nrn1 expression in bulk proteomic data from 5xFAD mice comparing control and PAP-1 treatment.
**Sheet#14(Suppl.Fig.4N):** Boxplot showing pseudo bulk Nrn1 expression in GABAergic neurons.
**Sheet#15(Suppl.Fig.4O):** Boxplot showing pseudo bulk Nrn1 expression in glutamatergic neurons.
**Sheet#16(Suppl.Fig.4P):** Boxplot of eigen protein expression for Mito Carta-associated proteins following Shk-223 treatment.
**Sheet#17(Suppl.Fig.4Q):** Expression of GFAP, SV2A, PSD-95, and TOMM20 in control and PAP-1 treated samples. Control and PAP-1 treated 5xFAD whole brain lysates have been compared using tandem mass tag (TMT)-based proteomics for different markers of gliosis(GFAP), synaptic integrity (SV2A, PSD95) and mitochondrial protein TOMM20.
**Sheet#18(Suppl.Fig.4R):** Statistical analysis of western blots remaining blots (STAT1 and STAT3) from **Fig5K** that compares control and PAP-1 5xFAD mice.

### Supp. Datasheet 5: Supp.Figure.5 analysis

**Sheet #1 (Suppl.Fig.5B):** Western blot and quantification of cell lysate of MMC upon Kv1.3 blockade using PAP-1 or Shk-223 followed by IFNα induction.
**Sheet #2 (Suppl. Fig. 5C)** Statistical analysis used in the graphs for expressional levels of different proteins from supplementary figure 5C

### Supp. Datasheet 6: Supp.Figure.6 analysis

**Sheet #1 (Suppl. Fig.6A)** Principal component analysis (PCA) of CSF proteomic data prior to normalization.
**Sheet #2 (Suppl. Fig.6B)** CSF proteomic dataset after removal of proteins with greater than % 75 missing values and subsequent imputation.
**Sheet #3 (Suppl. Fig.6C)** Principal component analysis of CSF proteomic data following normalization.
**Sheet #4 (Suppl. Fig.6D)** CSF proteomic abundance matrix following filtration, imputation and column sum median based normalization.
**Sheet #5 (Suppl. Fig.6E)** Variance partition analysis performed before correction for blood contamination.
**Sheet #6 (Suppl. Fig.6F)** variation of partition analysis was performed after correction of blood contamination.
**Sheet #7 (Suppl. Fig.6G)** Overlap of PAP-1 regulated proteins between brain and CSF proteomic datasets.
**Sheet #8 (Suppl. Fig.6H)** Overlap of differentially expressed proteins between brain and CSF comparing ShK-223 and PBS.
**Sheet #9 (Suppl. Fig.6I)** Correlation analysis of brain and CSF proteomic $log_2$ fold changes following PAP-1 treatment**.**
**Sheet #10 (Suppl. Fig.6J)** Correlation analysis of brain and CSF proteomic $log_2$ fold changes comparing ShK-223 and PBS.
